# 2500 Years of Human Betaherpesvirus 6A and 6B Evolution Revealed by Ancient DNA

**DOI:** 10.1101/2024.06.25.599715

**Authors:** Meriam Guellil, Lucy van Dorp, Lehti Saag, Owyn Beneker, Biancamaria Bonucci, Stefania Sasso, Tina Saupe, Anu Solnik, Helja Kabral, Raili Allmäe, Jessica Bates, Jenna M. Dittmar, Xiangyu Jack Ge, Sarah Inskip, Tõnno Jonuks, Victor N. Karmanov, Valeri I. Khartanovich, Maarten H. D. Larmuseau, Serena Aneli, Craig Cessford, Aivar Kriiska, Marika Mägi, Martin Malve, Natasja De Winter, Mait Metspalu, Luca Pagani, John E. Robb, Toomas Kivisild, Charlotte J. Houldcroft, Christiana L. Scheib, Kristiina Tambets

## Abstract

Human betaherpesviruses 6A and 6B are double-stranded DNA viruses, specialised in infecting humans and are best known as the main causative pathogens of the common childhood infection “sixth disease”. Despite only being discovered in the 1980s, these viruses are speculated to have a much longer and more complex history within the human population than available modern data make clear. The viruses are carried by large fractions of the human population and can integrate into the human genome, leading to a wide range of clinical manifestations of varied severity. Here, we present the first nine full and two partial ancient genomes of HHV-6A and 6B, dating as far back as the Italian Iron Age (ca. 1100–600 BCE). We demonstrate that large fractions of the current HHV-6 diversity were already well established in the human population by the 14th century CE. Our data suggests that HHV-6B integrated into the human genome at the latest before the 1st-6th century CE, with two integrated clades being populated by ancient DNA genomes in our phylogeny, which further supports that they originated from likely much older ancient founder events. Additionally, we show that all known inherited chromosomally integrated (ici-)HHV-6A clades were already represented in European historical populations, confirming that ici-HHV-6A no longer integrates into the human germline within populations of European ancestry and likely endogenized in early human history. Finally, our results demonstrate the unique suitability of archaeological remains and ancient DNA for the study of the evolution of integrated viruses in human populations.

## INTRODUCTION

*Human betaherpesviruses 6A* (HHV-6A) and *6B* (HHV-6B) are two distinct but closely related large double-stranded DNA viruses restricted to humans. Both viruses were first discovered in the 1980s from patients with lymphoproliferative disorders ^1,2^ and were initially classified as a single species ^3^. As such, these viruses are relatively understudied, with often unclear effects on human health and limited knowledge of their long-term co-evolution with the human host or the age of human association. Both HHV-6 viruses can cause mild or asymptomatic primary infections but are also closely associated with disease in immunocompromised individuals with complex disease courses. Whether the link of these conditions with HHV-6 is causal or merely an association remains to be determined. This uncertainty is likely due in part to the relatively recent discovery of the viruses and their ubiquity and lifelong latency within human populations, making it challenging to establish clear causal links. As such the viruses have been linked with a wide range of conditions such as transplantation-associated encephalitis ^4,5^, graft-versus-host disease ^6^, female infertility ^7,8^, myocarditis ^9,10^, multiple sclerosis ^11,12^ and preeclampsia ^13,14^ to name a few.

Today, HHV-6B infects around 90% of all children by the age of two, and is best known as the main causative pathogen behind *roseola infantum*, also known as sixth disease, the leading cause of febrile seizures in children, although primary infections have also been reported in adults ^15^. HHV-6A infections are epidemiologically and immunologically distinct from HHV-6B infections. However, the virus’s involvement in causing disease and its transmission modalities are less well understood. This is because, while HHV-6A is estimated to be phylogenetically older than HHV-6B, it is less prevalent in human populations today ^15,16^ and infections occur mostly in adulthood. A factor further complicating the study of the clinical impact of HHV-6A, is that due to the high prevalence of acquired HHV-6B infections, most individuals carrying HHV-6A would have been infected by HHV-6B at some point in their life ^17^.

Much like most herpesviruses, the HHV-6 viruses will remain latent in the infected host for life once acquired. However, contrary to all other known human herpesviruses, HHV-6 viruses achieve latency by integrating into the host genome. Latency can take place in a wide range of tissues and is detectable in saliva even after a primary infection following reactivation, which is the main mechanism of spread for *roseola infantum* ^15^. This usually happens in somatic cells, following an acquired exogenous infection, but rare integration events into germline cells, from where the virus can be transmitted vertically to ca. 50% of all offspring in a Mendelian fashion, have led to lineages of inherited carriage of HHV-6 sequences within every nucleated cell of the human body, which is estimated to be present in ca. 0.4–1% of the population worldwide ^15,18,19^. Integration sites typically occur between telomeres and subtelomeres and have been shown to impact the length and stability of telomeres in somatic integrations, with some hypothesising that shorter telomeres might be more frequent integration loci for HHV-6 ^15,19,20^. The consequences of the viral integration into telomeres are not well understood. However, the location of the integration could have a differential impact on the health of the host ^17^.

This wide range of latency mechanisms has led to markedly different evolutionary dynamics across clades within the virus’s phylogeny, some from acquired infections (acq) (circular/linear genomes, not integrated), non-inherited chromosomal integrations (ci) (linear genome, somatic) and inherited chromosomal integrations (ici) (linear genome, germline) ^17^. While acquired infections evolve similarly to non-integrating infecting viruses (engage in recombination with typically higher mutation rates), integrated versions of the virus show lower genetic diversity in clades, probably due to the higher fidelity achieved by human genomic replication. Additionally, integrated versions of the virus are capable of (re)activation, leading to the release of virions (circular genome) ^17,21^.

While the seroprevalence of HHV-6B is estimated to be over 90% in modern populations, this number is hard to verify in ancient and historical populations due to the fact that the most common form of latent carriage would be limited to a small number of somatic cell integrations as opposed to ubiquitous germline inherited integrations. This makes detection of acq/ciHHV-6B in hard tissue, which is the most common form of tissue available for ancient DNA (aDNA) research, likely difficult. On the other hand, germline chromosomal integrations are ideal for aDNA detection, particularly in samples with high amounts of host DNA. However, inherited infections are much rarer and only present in a small percentage of the human population today ^15^. Further, these carriage rates could have been much lower in human population the closer to the initial integration events they might date, as they are speculated to originate from a small number of human ancestors ^22^. However, modern integrations have also been described for HHV-6B and no direct data is available to verify the age of the integrated clades described in the literature ^16^ and it remains an open question whether HHV-6A is still capable of integrating into the human germline. The timing of such ancient integration events is still unclear but Aswad et al. ^16^ estimated that they occurred more than 50,000 years ago, which would place it around or before the time of the major human migration out of Africa.

Human herpesviruses have been reported in ancient samples (*Herpes simplex virus 1* ^23^ and Epstein–Barr virus ^24^), but full genomes have only been made available for *Herpes simplex virus 1*, where the addition of dated aDNA genomes enabled a more accurate estimation of the emergence dates of the viral pathogen. This in fact is one of the undisputable advantages of aDNA for the study of dsDNA/ssDNA viruses, which, comparatively to RNA viruses, evolve at much slower rate and therefore can be challenging to study in the context of long-term evolutionary dynamics using solely modern data. Additionally, integrated viral sequences remain largely unstudied in the historical population, despite the unique opportunity given by aDNA datasets to study the early evolution of endogenous viruses and potentially the early phase of their endogenization in the human germline. The only herpesvirus, which is known to integrate into the host genome, that has been recovered from aDNA to date is the *Gallid alphaherpesvirus 2*, which causes Marek’s disease in chicken ^25^. Here, we describe the first eleven ancient genomes of two integrating viral pathogens, HHV-6A and HHV-6B, and trace 2500 years of the viruses’ evolution in the European population.

## RESULTS

### Identification and authentication of HHV-6A and 6B

During the metagenomic screening of larger aDNA population genetics projects, we recovered sequences matching HHV-6 species from a total of eleven human samples spanning from the 8th-6th century BCE to the 14th century CE (see Fig. 1). Of these, six samples were identified as carrying HHV-6A and five as carrying HHV-6B, through taxonomic classification and comparative mappings (see Table S9). Differentiation from the closely related *Human betaherpesvirus 7* (HHV-7) was further validated by the presence of the U83 and U94 genes, which are specific to HHV-6 species and suspected to be involved in the establishment of latency ^15^. Deamination signatures and fragment length (see Table S2 and Fig. S7-S9) were consistent with ancient DNA.

**Figure 1:**
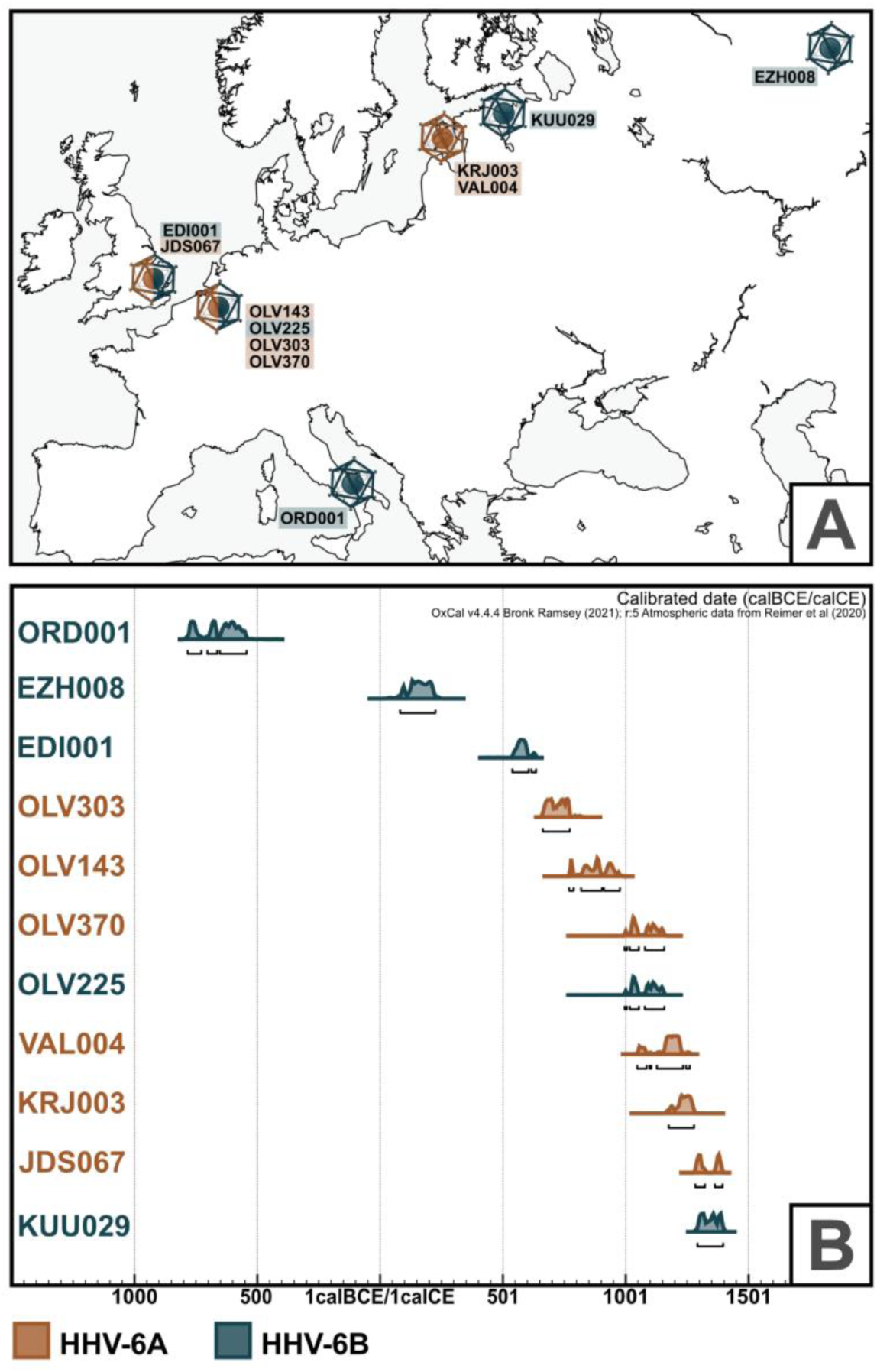
A) Map of Europe depicting all sampling locations and Human betaherpesvirus 6 species found at the site. B) OxCal plot with calibrated radiocarbon dates for all samples presented in this study. Dates are shown in calBCE/calCE (EZH008 dating is likely affected by the freshwater reservoir effect see SI).

### Wide temporal, geographic and cultural range of identified individuals carrying the virus

The samples analysed in this study stem from a diverse group of individuals in terms of age-at-death, genetic sex and geographic provenance (see Table 1). Sample ORD001 stems from a female individual from Iron Age Italy, EDI001 from a male aged 16–18 years from the Anglo-Saxon site of Edix Hill (England), and JDS067 from a male child excavated at the Hospital of St. John the Evangelist in Cambridge (England). Four of our samples, three female adults and one male adult, originated from a medieval parish graveyard in Sint-Truiden (Belgium). An adult male, an adult woman and an infant girl were recovered from medieval Estonian sites, and finally, an older male originated from a site in Ezhol (Komi Republic, Russia). This diversity is further reflected in their cultural and temporal spectrum (see Fig.1 and SI for additional archaeological and osteological data, as well as Table S10 for expanded radiocarbon dating data), with dates ranging between the 8th-6th century BCE and the 14th century CE. While all other HHV-6 positive individuals belonged to farming societies, the individual from Ezhol demonstrates that HHV-6B was also transmitted among taiga hunter-fisher-gatherers. The only site for which more than one individual carrying the viruses could be detected is Sint-Truiden, but only two individuals (OLV225 and OLV370) can be considered contemporaneous.

**Table 1:** Sample metadata and mapping statistics for all new ancient DNA samples described in this study. Mapping statistics include the mean depth of coverage, the mean edit distance (MQ>30) and the percent of sequence coverage at depths of coverage one, three, and five. Further, we describe the percentage of endogenous human DNA, based on our mappings to the human genome, and expected integration sites for the recovered viral genomes based on our phylogenetic analysis (Fig. 3 & 4), and on Aswad et al ^16^. An expanded version of this table is available in Table S1. Calibrated C14 dates are rounded. OxCal calibration plots are available in Fig. S5 and S6, and uncalibrated C14 dates in Table S10. Full mapping statistics are available in Table S2.

| Sample ID | Archaeological Site | Geographical Location | Sample Type | Rounded Cal. C14 Dates | Age | Genetic Sex | Species Found | Mean Depth of Coverage | Mean Edit Distance (MQ>=30) | % 1X | % 3X | % 5X | 3p C>T (Pos 1) | Expected Integration Site | Endogenous Human DNA % |
| --- | --- | --- | --- | --- | --- | --- | --- | --- | --- | --- | --- | --- | --- | --- | --- |
| utigEDI001 | Edix Hill | England, Cambridshire | Tooth Root , Bone | 545-640 cal CE | 16-18 years | XY | HHV-6B | 16.59 | 0.82 | 99.3 | 93.4 | 90.2 | 22% | 19q | 46.59 |
| utigEZH008 | Ezhol | Russia, Kharino | Incus Bone | 75-220 cal CE | ca .50 years | XY | HHV-6B | 8.11 | 0.62 | 97.8 | 80.1 | 70.9 | 26% | 17p | 75.42 |
| utigJDS067 | St Johns | England, Cambridshire | Tooth Root | 1280-1390 cal CE | 9-10 years | XY | HHV-6A | 3.42 | 1.14 | 84.7 | 32.5 | 20.5 | 22% | 17p | 3.72 |
| utigKRJ003 | Karja cemetery | Estonia, Saaremaa | Tooth Root | 1175-1275 cal CE | 30-35 years | XY | HHV-6A | 10.33 | 1.11 | 96.5 | 90.3 | 86.7 | 14% | 18q | 24.03 |
| utigKUU029 | Kukruse | Estonia, Ida-Viru | Petrous Bone | 1300-1400 cal CE | 10.5-18 months | XX | HHV-6B | 1.85 | 0.49 | 43.4 | 1.8 | 1.3 | 13% | NA | 83.33 |
| utigOLV143 | Sint-Truiden | Belgium, Limburg | Tooth Root | 770-975 cal CE | Adult | XX | HHV-6A | 10.05 | 0.98 | 96.9 | 89.8 | 85.8 | 16% | 18q | 78.04 |
| utigOLV225 | Sint-Truiden | Belgium, Limburg | Tooth Root | 995-1155 cal CE | 20+ years | XX | HHV-6B | 26.56 | 0.51 | 99.6 | 98.6 | 98.2 | 12% | NA | 83.07 |
| utigOLV303 | Sint-Truiden | Belgium, Limburg | Tooth Root | 665-775 cal CE | 20-40 years | XX | HHV-6A | 6.76 | 1.05 | 95.4 | 78.1 | 67.2 | 14% | 19p | 69.69 |
| utigOLV370 | Sint-Truiden | Belgium, Limburg | Tooth Root | 995-1155 cal CE | 20-40 years | XY | HHV-6A | 6.80 | 1.00 | 95.4 | 77.4 | 65.7 | 15% | 18q | 23.13 |
| utigORD001 | Ordona | Italy, Apulia, Foggia | Petrous Bone | 780-540 cal BCE | 9-11 years | XX | HHV-6B | 1.77 | 1.59 | 40.1 | 4.6 | 3.2 | 10% | NA | 4.02 |
| utigVAL004 | Valjala Church Yard | Estonia, Saaremaa | Tooth Root | 1045-1260 cal CE | 40-50 years | XX | HHV-6A | 11.39 | 1.14 | 96.9 | 91.6 | 89.3 | 16% | 18q | 68.27 |

### Host DNA analysis reveals high endogenous DNA content in most samples

The endogenous human DNA content was below 5% for JDS067 and ORD001, but above 20% for all other samples and above 65% for six samples (see Table S7). The average genomic coverage was below 0.1X for three samples (ORD001, JDS067, KRJ003), above 1X for another four samples (OLV143, OLV303, OLV225 and EDI001), and between the two for the rest. We scanned our mappings for clinical variants of interest in the context of an HHV-6 carriage but did not detect any clear occurrence of such (see SI and Table S8).

### Tissue tropism and sampling significance for HHV-6 carriage type

For this study, the tissues and anatomical sites from which the viral sequences were recovered are highly relevant, as this could evidence the type of viral carriage recovered from each host. For JDS067, VAL004, KRJ003 and all OLV individuals, the only available sample types were teeth (see Table S1), allowing for the possibility of both acquired and inherited infections. However, for ORD001 and KUU029 HHV-6 sequences were recovered from petrous bones and for EZH008 from an incus bone. These bones are very dense with low remodelling potential and vascularization, making an acquired active infection or a localised somatic integration or latency rather unlikely. Instead, these sampling sites point to the probability of inherited carriage of the viruses. EDI001 was a singular case for which multiple sample types were available (tooth, bone), and the virus could be detected in all sample types, making a clear case for an inherited germline integration in this individual.

### Comparative analysis and taxonomic classification show clear species identifications

Upon identification of HHV6 sequences in screening shotgun datasets and enrichment (see methods), all datasets were mapped to the HHV-6A (NC_001664.4) and HHV-6B (NC_000898.2) reference sequences (ca. 90% sequence identity). Additional mappings to the phylogenetically closest herpesvirus, *Human betaherpesvirus 7* (NC_001716.2), were also performed to exclude potential misidentifications (see SI), of which there were none. Based on the mapping statistics and coverage, samples were classified into HHV-6A and HHV-6B samples, which coincided in all cases with the original taxonomic classification using KrakenUniq ^26^ (see Table 1).

### Phylogenetic analysis yields termini ante quem for iciHHV-6 clades

Of the eleven ancient HHV6 genomes recovered, nine achieved depths of coverage high enough to generate high quality SNP calls and perform a phylogenetic analysis (see Table S2). The lowest coverage sample, for which we still recovered a full genome, JDS067, had a mean depth of coverage of 3.42X. All other full genome samples were above 5X (between 6.76X and 26.56X) (see Table 1 and Fig. 2). Partial genomes, under 3X, were not used for phylogenetic analysis (KUU029, ORD001). Following reference mapping for HHV-6A and HHV-6B respectively, SNPs were called using freebayes ^27^ and consensus sequences were produced using bcftools ^28^ while masking for terminal repeats and repetitive/low complexity intervals (see SI). 278 modern genomes (see Table S3) from NCBI were used for the analysis ^16,22,29–40^. The concatenated alignments were checked in RDP5 ^41^ using the phi-test, which revealed “very good evidence for recombination”, as expected for HHV-6 viruses. Recombinant intervals were then identified and masked in the alignment using RDP5 (see SI). Informative sites at 95% deletion were kept, resulting in an alignment length of 1,408 bp for HHV-6A (n=72) and 1,942 bp for HHV-6B (n=217). Maximum likelihood phylogenies were generated with IQ-Tree2 ^42^ using ultrafast bootstrap and SH-aLRT support calculations with 10,000 replicates each. IQ-Tree2’s integrated ModelFinder selected the TVM+F+ASC and TVM+F+ASC+R3 models for HHV-6A and HHV-6B respectively.

**Figure 2:**
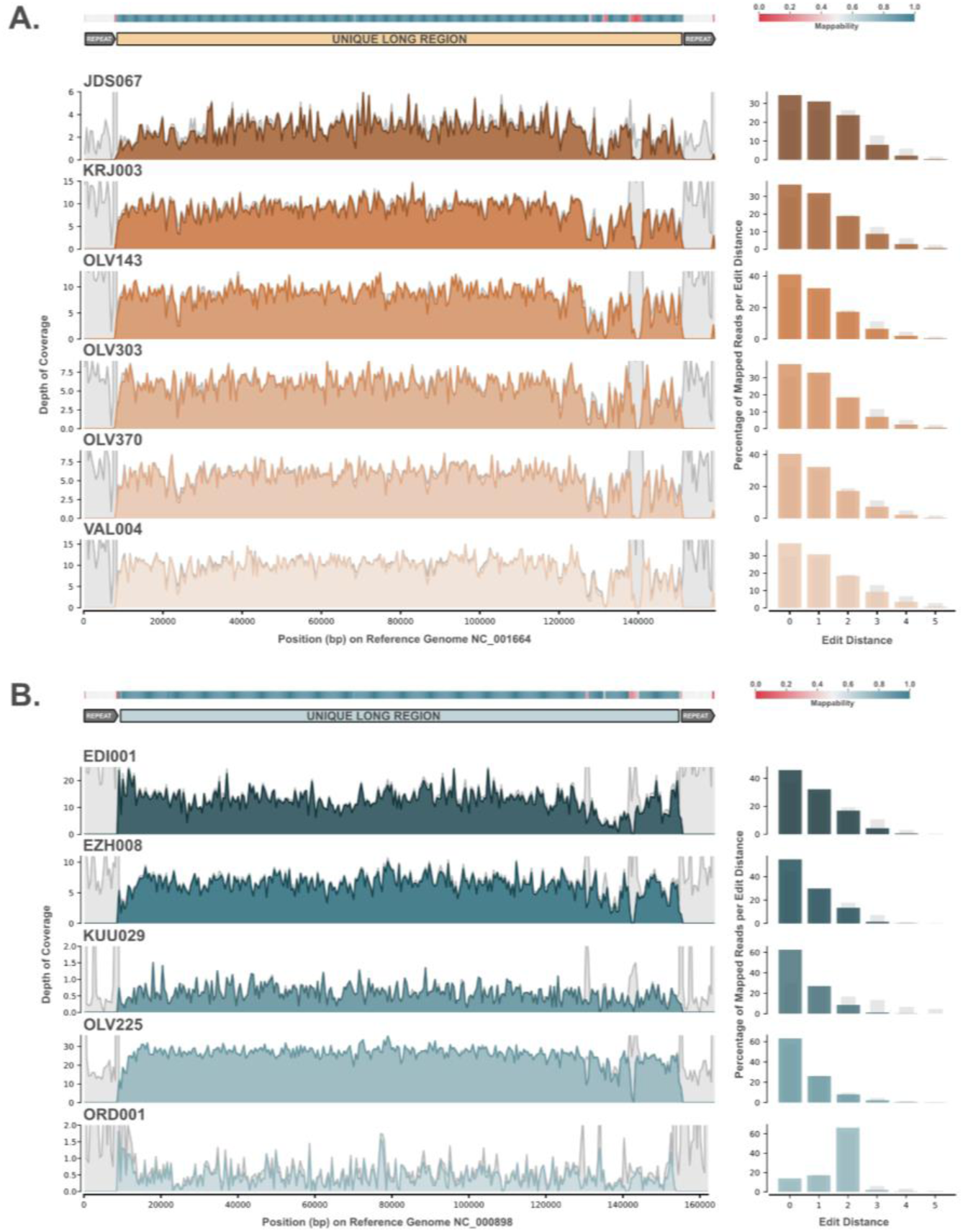
Left: Genomic coverage plots for all A) HHV-6A and B) HHV-6B mappings to respective reference sequences. Intervals with reads under a mapping quality of 30 are shown in light grey. On the top of each figure, mappability of the reference sequence is plotted in a heatmap and the second line shows the location of the unique long region of the 6A and 6B reference genomes used for analysis. On the right, edit distances for the mappings are shown in a barplot with light grey bars showing the percentage of reads under a mapping quality of 30.

Overall, the phylogenetic position of all ancient genomes is well-supported and clearly nested within known integration clades (see Fig. 3 and 4), which show distinct evolutionary dynamics due to their replication within the human genome and therefore showcase extremely low mutation and recombination rates when compared to non-vertically inherited viruses. These clades can also give us insights into the likely integration site for each strain ^16^. An exception is sample OLV225, which is not nested in an extant defined clade associated with known human genome integration loci. This genome clusters with two ci/ici genomes from relatives from the USA with likely European ancestry (HP40E6 and HP43E10), which could hint at undersampled diversity in this lineage. However, this clade, situated basally on the branch giving rise to the B5 and B8 clades, is less well-supported than the integrated clades and more sensitive to pruning during recombination analysis. While integration cannot be excluded for this sample due to relatively high host DNA content in the datasets, the sample type (tooth) and the strain’s phylogenetic placement do not irrevocably point towards an ici or ci strain. Contrary to Aswad et al, sample NY-390 does not cluster with the B5 clade, which seems in line with it being an acquired strain and of hispanic ancestry, and thus phylogenetically distinct from the integrated B5 clade. In our analysis, clade B1 was further split into two subgroups: one exclusively populated by Asian genomes and a second with non-Asian genomes, which is closer to the main B7 clade. Some highly divergent genomes, such as NY-310, were omitted from the main phylogeny to avoid reducing the analysable sequence further.

**Figure 3:**
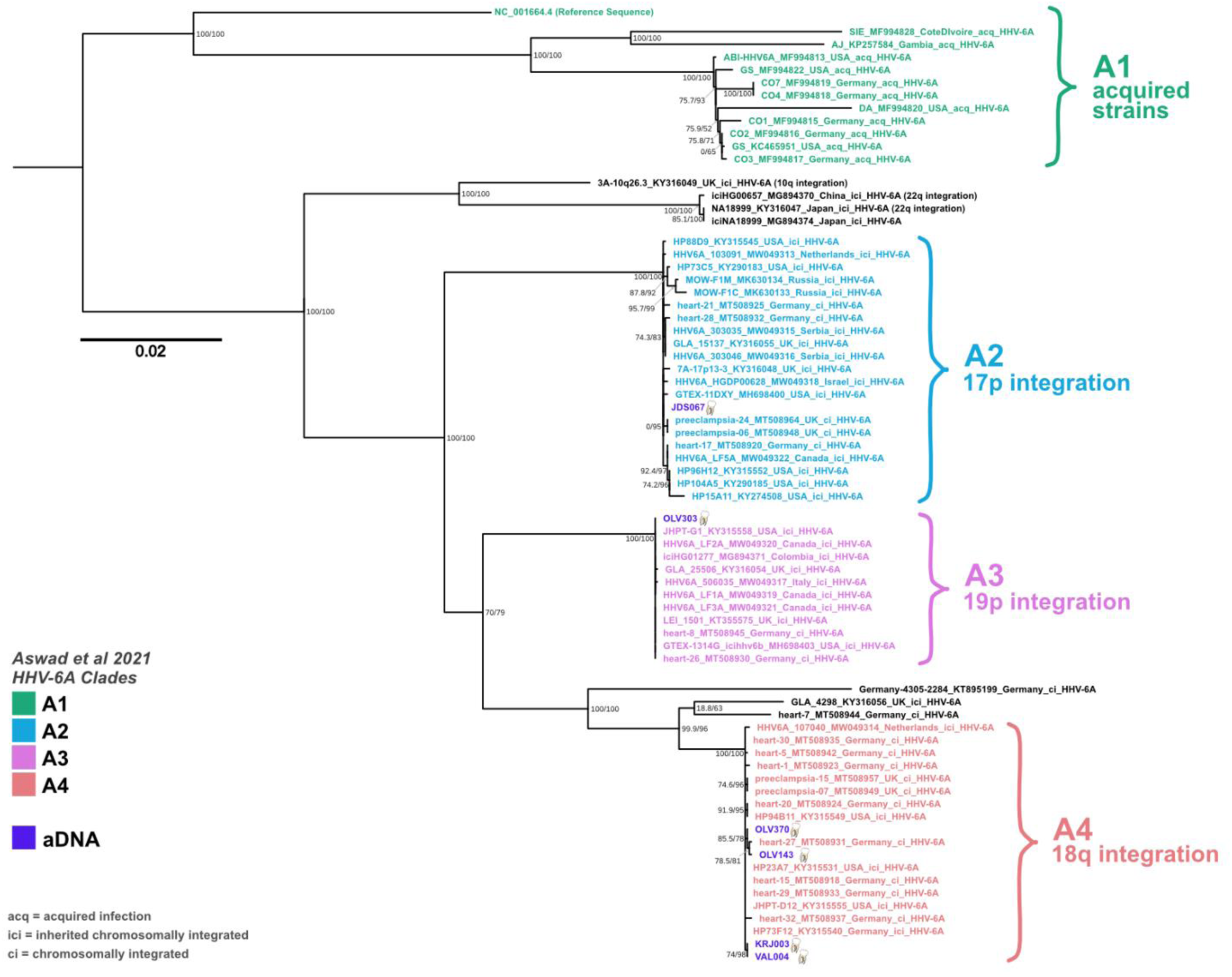
Midpoint-rooted maximum likelihood phylogeny for Human betaherpesvirus 6A using 66 modern and 6 ancient genomes. Node labels show SH-aLRT support (%) / ultrafast bootstrap support (%) based on 10,000 replicates each in IQ-Tree2. Tip labels are coloured based on clade groupings and integration locations are shown as defined in Aswad et al. 2021. Ancient DNA samples are shown in purple, with illustrations of the type of samples the genomes were extracted from, either with a tooth or a petrous bone (also for incus bone). Strains are only noted as ici, if they are known to be inherited.

**Figure 4:**
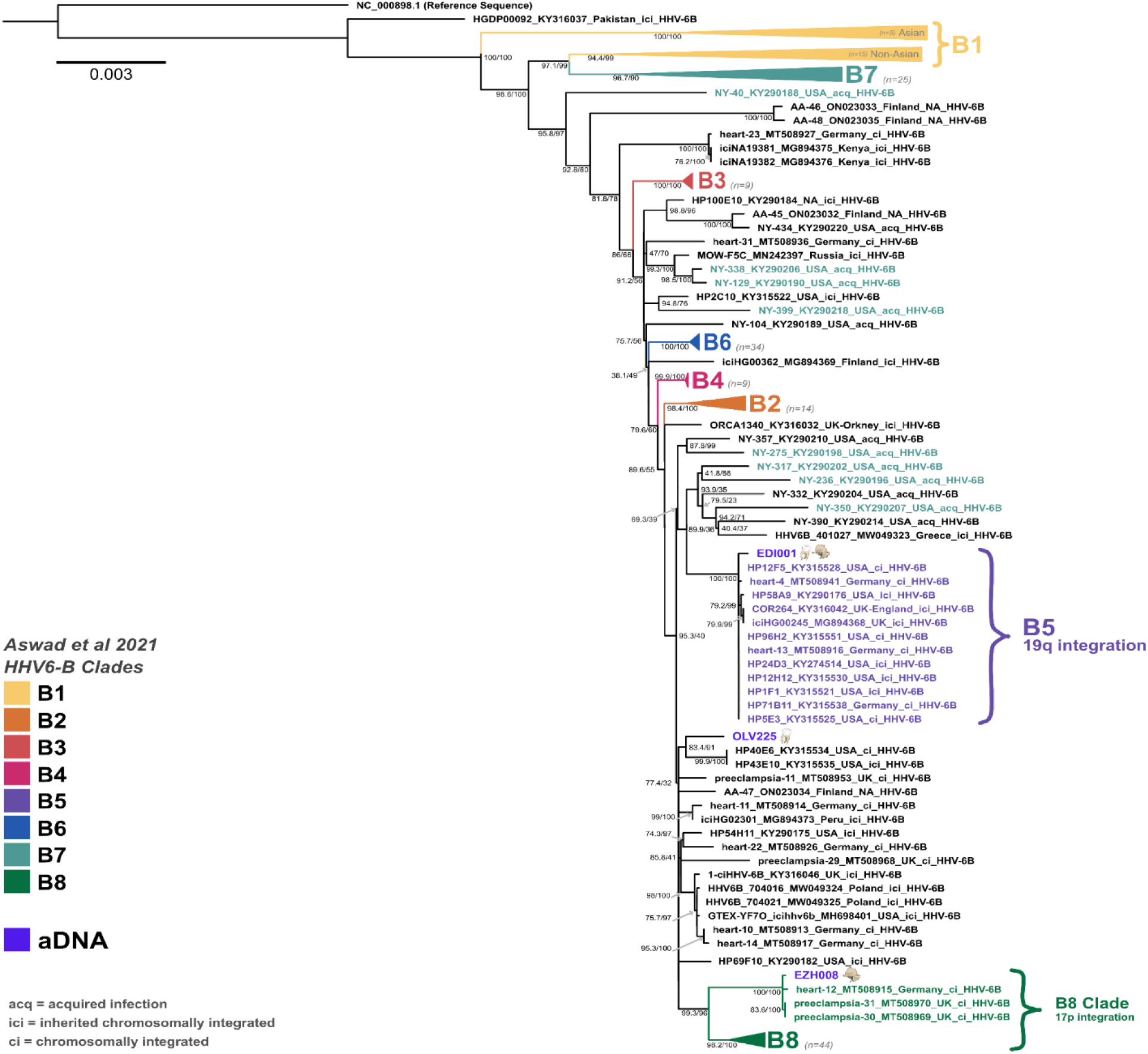
Midpoint-rooted maximum likelihood phylogeny for Human betaherpesvirus 6B using 214 modern and 3 ancient genomes. Node labels show SH-aLRT support (%) / ultrafast bootstrap support (%) based on 10,000 replicates each in IQ-Tree2. Tip labels are coloured based on clade groupings and integration locations are shown as defined in Aswad et al. 2021. Ancient DNA samples are shown in purple, with illustrations of the type of samples the genomes were extracted from, either with a tooth or a petrous bone (also for incus bone). Strains are only noted as ici, if they are known to be inherited.

Based on our results, we can observe that all integrated HHV-6A clades are represented in the European population by the 14th century CE at the latest, but likely have a much older evolutionary history. Clade A2 is represented by one English genome dating to the 13th–14th century CE, clade A3 by one Belgian genome dating to 7th–8th century CE and clade A4 is represented by four genomes from Estonia and Belgium dating between the 8th and the 13th century CE. For HHV-6B, our phylogeny yielded *termini ante quem* of the 1st– 6th and 6th–7th century CE for clade B8 and B5 respectively. While we could only detect HHV-6A genomes in individuals who died after the 7th-8th century CE, which is in contrast to the assumption that the HHV-6A species is older than the HHV-6B species, this is likely due to sampling biases rather than to unexpected emergence dates, which will likely be resolved by the addition of more ancient genomes in the future.

### Lack of global detectable temporal structure

We additionally set out to ascertain whether we could recover meaningful temporal information in the alignments through the inclusion of ancient genomic observations of HHV-6A and HHV-6B. Through consideration of topologies of mixed evolutionary behaviour (concurrently considering acquired and integrated clades) and those expected to exhibit a consistent evolutionary behaviour (acquired only or by integrated clade) we were consistently unable to recover a robust pattern of temporal evolution (see SI). The exception was the HHV-6A clade of A2 integrated viruses where we were able to obtain a temporal correlation, though noting this was predominantly driven by the placement of JDS067, our lowest coverage full genome, which was itself of uncertain topological position, either slightly basal to the A2 clade or within it (see Fig. 3 and S15-17), depending on the analysis and length of the final alignment. Hence, we did not proceed to formal tip-calibration approaches. While not able to leverage phylogenetic information for estimation of divergence times, a major advantage of the inclusion of aDNA samples is that it allows the establishment of the absolute minimal age of a clade based on sample associated radiocarbon information.

### Likely iciHHV-6B carrier probably succumbed to the plague

From the eleven individuals in which we could detect HHV-6 sequences, EDI001 is known to have been infected by another pathogen. A plague infection had previously been reported for this individual in Keller et al. 2019 ^43^. The sample stems from the site of Edix Hill, United Kingdom, which is well known for its plague cases ^43^, dating back to the First Plague Pandemic, and has yielded co-infections in the past ^44^. As we expect EDI001 to have carried an inherited chromosomally integrated HHV-6B strain, it is unlikely to have been involved in the death of the individual, as they likely died of an *Yersinia pestis* infection which kills rapidly.

## DISCUSSION

The HHV-6 viruses are a relatively recent addition to our knowledge of herpesviruses and endogenous viruses overall. Hence, we are only beginning to piece together how and for how long they have impacted human health. Some symptomatic presentations, such as the common childhood disease *roseola infantum*, are better understood, but the impact of integration into human genomes (somatic or germline), and particularly the transmission of integrated variants via the germline, are still hard to elucidate. An additional by-product of their novelty in research is that no samples pre-dating the 1980s are available, thus reducing the temporal range and signal available for evolutionary estimates even more.

### Ancient DNA can shed light on the evolution of HHV-6A and 6B

Ancient DNA provides the unique opportunity to bridge this gap and study DNA viruses over their history as they evolved within and with human populations (e.g. Herpes simplex virus 1 ^23^, Hepatitis B ^45^ and Variola ^46^). In this study, we reconstruct nine full and two partial *Human betaherpesvirus 6* genomes from individuals dating between the 8th-6th century BCE to the 14th century CE, expanding our knowledge to more than 2500 years of these viruses’ evolutionary histories. The genomes originate from individuals of all ages and both sexes, which were recovered from across Europe. Of these, we identify at least nine aDNA genomes to be likely iciHHV-6 genomes. For EDI001 (England, 545-640 cal CE) we were able to detect HHV-6B across multiple tissue types and the sample clustered within an endogenous clade (B5) known to integrate into Chr19q. EZH008 (Russia, 1st-6th century CE), our second-oldest sample, clustered in an endogenous sub-clade (B8) known to integrate into Chr17p and was isolated from an incus bone. JDS067 (England, 1280–1390 cal CE), KRJ003 (Estonia, 1175-1275 cal CE) and VAL004 (Estonia, 1045–1260 cal CE) were all isolated from teeth but also clustered in endogenous clades with known HHV-6A integration loci (Chr17p for JDS067 and Chr18q for the Estonian samples).

Samples from Sint-Truiden (OLV143, OLV225, OLV303, OLV370) stem from a large population genetics project (in prep ^47^) involving extensive sequencing (130 million raw sequence reads, on average, generated for 404 individual samples), reducing, while not removing, the likelihood of large undetected groups carrying HHV-6 integrations within the population. However, based on radiocarbon dates (see Fig 1 & Table 1) only two individuals, OLV370 and OLV225, are likely to have been contemporaneous. Nevertheless, the site confirms the concurrent presence of both HHV-6A and HHV-6B within the same population around the 10th– 12th century CE in Europe. They were all isolated from teeth. OLV143 and OLV370 cluster in the same clade as VAL004 and KRJ003 (A4). OLV303 is the only representative of the endogenous clade A3 (Chr19p) and OLV225 is the only HHV-6B sample recovered at the site and the only full genome which cannot irrevocably be placed in a known endogenous clade.

### Partial genomes can still inform us about HHV-6 endogenicity

Coverage across the human genome and the viral genome cannot be directly correlated in this study, as a) the first stems from shotgun sequencing data and the latter from enriched genomic libraries and b) HHV-6 viruses integrate into telomeric regions, which can complicate sequencing efforts due to variation in telomere stability/length and sequence complexity around the integration site. However, for all our samples one trend remained clear: samples for which high coverage of HHV-6 genomes could be achieved and which clustered in integrated clades also showcased high levels of host DNA preservation (22.7–83.07%) and vice versa. As we expect most of our genomes to be from germline integrated lineages, this observed trend coincides with expected observations and persists whether genomes were isolated from petrous bones, incus bones or teeth.

KUU029 was the exception to this trend. The mean depth of coverage for sample KUU029 (Estonia, 1300– 1400 cal CE) remained low, even following target enrichment (see Table 1); this sample is of interest as it is likely the only acquired or somatically integrated strain in our sample set, despite having been recovered from a petrous bone. Although endogenous host DNA is estimated at 83.33% for this sample, the mean depth of coverage for our mapping against the HHV-6B reference sequence is 1.88X after target enrichment. Further, the sample stems from an infant (10.5–18 months old), which makes the likelihood of an acquired infection higher, as today *roseola infantum* is the most common active infection caused by HHV-6B ^48^ and usually manifests in children between the ages of 6 months and two years. The reason for the recovery of an acquired or recently somatically integrated virus from a petrous bone could be the age of the individual, as their skull would have still been in the early stages of cranial development.

### The oldest and most divergent genome: ORD001

The second sample, ORD001 (Italy, 780–540 cal BCE) has the lowest depth of coverage of all samples, but is also the oldest sample in our analysis, dating to the Italian Iron Age (ca. 1100–600 BCE). Our mapping shows a wider divergence from the HHV-6A and HHV-6B reference sequences (mean edit distance 1.59), while still being assignable to HHV-6B based on our analysis (see SI and Figure S13 and Table S9). This is in stark contrast to the other HHV-6B samples for which the mean edit distance never exceeded 0.82. This sample likely diverged significantly from the reference sequence and showcases how dynamic and convoluted we can expect HHV-6B evolution to have been even in prehistory. This could either point to the presence of a very basal divergent form of HHV-6B or an ancestral form of HHV-6B. However, more sequencing efforts, from both modern and ancient DNA, are needed to determine whether an ancestral form of HHV-6B might have been circulating in the European Iron Age, as older strains and closely related modern comparative sequences (e.g. from non-human primates), are still missing.

### Geographical structure across the aDNA HHV6 genomes

Geographical clustering in our phylogenies is limited within the available aDNA genomes, which could change with more data. Samples KRJ003 and VAL004 show clear geographical clustering, which is not surprising considering they stem from the Island of Saaremaa in the Baltic Sea (Estonia) and are of similar age (12th-13th and 11th-13th century CE). The same integrated clade (A4), also harbours two genomes from the Belgian site of Sint-Truiden (OLV143, OLV370) dating to the 8th and 10th, and to the 10th to 12th century CE, respectively. These groupings clearly show that clade A4 was already well dispersed in Europe and was even established in the rather remote Saaremaa population by the 11th-13th century CE. All other high-coverage genomes (EDI001, EZH008, JDS067, OLV225, OLV303) cluster in separate clades on their own, highlighting the heterogeneity of our sample set.

### HHV-6A no longer integrates into the genomes of populations with European Ancestry

The HHV-6A phylogeny is clearly separated into circulating and endogenous genomes, with no known overlap and long branches separating each clade. One of the big open questions regarding the evolution of HHV-6A is whether the virus still integrates into the human genome or if all integration events are ancestral ^16,19^. Contrary to HHV-6B where the evolutionary line between circulating and endogenous viruses is much more blurry, host-pathogen evolution might have impacted its ability to integrate into the germline sometime in human history ^16^. Based on our ancient data, we can say that all known main endogenous clades (A2, A3 and A4) were already represented in historical populations. This shows that the sampled diversity found within populations of European ancestry today is in all likelihood limited to ancestral integration events, and that these events are no longer occurring in European populations or only occurring in rare, undetected events. While we do not have much data for Asian strains, a small seemingly monophyletic clade of iciHHV-6A from eastern Asia is known to integrate into Chr22q and based on prior studies is also likely to stem from a single ancestral founder event, further validating this trend for the species ^19,49^.

Overall, the current HHV-6A phylogeny suffers from a heavy geographical sampling bias, with most of its data stemming from individuals with European ancestry, making any assumption on a global level difficult. This is further exacerbated by varying carriage rates for iciHHV-6A within populations ^19^, which are also generally lower than for iciHHV-6B (they make up only 10-40% of iciHHV-6 ^17^). However, in our case, sampled ancient and modern diversity overlap, and we can see a clear temporal continuity with ancient integration events leading to eventual long-term endogenization of the virus without reaching fixation, which were all in place at the latest between the 7th and 14th century CE. We can therefore confirm that endogenous and circulating HHV-6A genomes diverged in early human history, with clades A2-A4 originating from ancestral founder events into the human germline and that these integration events occurred in Europe prior to the colonial era.

### The evolution of HHV-6B was already convoluted in the human past

The evolutionary history of HHV-6B is much more convoluted and prior studies have demonstrated that endogenous strains of the virus likely serve as an endemic reservoir for horizontal transmission and integration events that likely still occur to this day ^16,21^. This means that endogenous and circulating strains are more closely related and interspersed across the phylogeny than for HHV-6A. Our ancient HHV-6B genomes cluster in integrated clades B5 and B8. All genomes for which the donor ancestry is known in clade B5 are likely to be of European ancestry. Clade B8 is also predominantly populated with genomes stemming from donors of likely European ancestry, with one exception being HP46B12, for which African ancestry is assumed based on previously published data ^35^. The last ancient HHV-6B strain clusters near two known endogenous genomes (HP40E6/HP43E10), which stem from first-degree relatives from North America with likely European ancestry ^16,29^. While it could theoretically hint at the presence of an undersampled integrated clade, this grouping is not as well-supported as other integrated clades, experienced more pervasive recombination and shows higher sequence divergence when compared to other integrated clades. The sample is likely nested within a complex basal branch, giving rise to the clades B5 and B8 (both populated by aDNA samples), the HP40E6/HP43E10 cluster and an amalgamation of acquired and endogenous strains. It can be assumed that the clades B5 and B8 originated from ancestral integration events much like all main iciHHV-6A clades, with our earliest *terminus ante quem* for iciHHV-6B dated to the 1st–6th century CE and our oldest HHV-6B sample dating to the 8th–6th century BCE. While we were unable to formally date the age of integration of clades, even upon inclusion of ancient observations, we anticipate it is likely that, with additional data to support a within clade sampling time-series or the detection of much older observations, this may be a feasible approach to assess long-term evolutionary rates in the viral species.

### How can we study HHV-6 viruses using ancient DNA?

While HHV-6 viruses are ubiquitous today, they mostly remain latent somatically and are thus much harder to detect and reconstruct using hard tissue samples, which are the most common sample types for aDNA analysis. On the other hand, iciHHV-6 would theoretically be much better suited for detection in aDNA datasets, as we can expect to find the virus in every human cell from birth and high viral loads in all tissue types. However, chromosomally integrated carriage is much less prevalent, with only an estimated 0.4–1% of the human population being carriers today ^15,19^. This percentage can be estimated to grow even smaller the closer we come temporally to the endogenization of the virus into the human germline.

Based on our data and the latency mechanism of the virus, we conclude that aDNA detection of HHV-6 viruses is indeed most likely to occur in the case of chromosomally integrated variants, as most of our recovered genomes cluster in endogenous clades or are likely to be integrated, despite iciHHV-6 genome carriage being much rarer than wild-type infections. This means that aDNA datasets will in all likelihood be biased towards the retrieval of iciHHV-6 but not restricted to it, as shown by sample KUU029. The reconstruction of aDNA HHV-6 sequences is facilitated by the fact that it can be retrieved from high endogenous DNA samples with relative ease, as it is part of the human genome and replicates with it. This is also the case in sample types which are only rarely thoroughly metagenomically analysed, which is exemplified by samples EDI001 (maxilla), KUU029/ORD001 (petrous bone) and EZH008 (incus bone). A second possibility would be the recovery of viral sequences originating from a somatic integration, e.g. into the salivary glands, from where it could have reactivated and shed into the oral cavity, as HHV-6B is known to be detectable in the saliva of healthy individuals well after their primary infection ^15^. However, HHV-6 might still often remain undetected due to bad overall DNA preservation, and their tendency to integrate into short telomeres might further impact their retrieval. Nevertheless, this study demonstrated that aDNA is a powerful tool for studying HHV-6 endogenization over the entirety of human history.

## CONCLUSION

Our data shows that HHV-6A & 6B were already well established in human populations prior to the 14th century CE, with our oldest genome ORD001 dating to the Italian Iron Age, around 8th–6th century BCE, clearly showcasing the long evolutionary history of these endogenous viruses in the European populations. We provide absolute minimal ages for all defined endogenous iciHHV-6A clades and conclude that the virus is no longer integrating into the germline in populations of European ancestry. Further, we demonstrate that HHV-6B was already present in the European Iron Age. We find that evolutionary dynamics seemed to have remained unchanged at least from the 1st-6th century CE until today, with HHV-6B having a more convoluted evolution without a distinct line between circulating and endogenous diversity as opposed to HHV-6A, for which aDNA genomes are distinctly segregated by integration loci. Finally, this study clearly demonstrates the power of aDNA research to study the evolution of HHV-6 viruses and other endogenous viral elements. More data, particularly from early human history but also from a wider geographic range will be key to further our understanding of the emergence and the endogenization of these viruses, which we now know have accompanied us since at least the Iron Age but are still novel to research today.

## Acknowledgments

We would like to thank Mari Tõrv for offering feedback on this project, as well as her knowledge of previous work on the site of Kukruse. We would also like to thank Benedikt Kaufer and Dirk Lassner for providing sampling dates and information for the heart cohort genomes used for our analysis. Analyses were carried out using the facilities of the High-Performance Computing Center of the University of Tartu. We would also like to thank the Cambridgeshire County Council. This work was funded and supported by: the Estonian Research Foundation Grant PRG1027 (to K.T., M.G., T.K., S.S., M.Ma. and L.S.), the Austrian Science Fund FWF FW547001/ESP162-B (to M.G.), the Estonian Ministry of Education and Research TK215 (K.T., T.K., L.S., S.S., B.B., A.K., M.T., M.Ma., H.K.), the FWO Grant G0A4521N (T.K., O.B., M.H.D.L., C.L.S., K.T.), Wellcome Trust Award no. 2000368/Z/15/Z (C.L.S., S.A.I., J.E.R., T.K. and C.C.), St John’s College University of Cambridge ( to C.L.S.), the Department of Genetics University of Cambridge (to C.J.H.) and the Foundation Osiliana (to M.Mä.).

## Authors Contributions

Study design and conception by M.G.; Funding Acquisition by K.T., M.Me., T.K., J.E.R., C.L.S. and L.P.; Laboratory work was performed by M.G., O.B., C.L.S., B.B., S.S., T.S., H.K., A.S., J.B. and X.J.G.; Analysis of the HHV-6 data was done by M.G. and C.J.H.; Temporal analysis was performed by L.v.D.; Analysis of the human data was performed by L.S., M.G. and C.J.H.; Archaeological and osteological background and data were provided by J.M.D., S.I., T.J., V.N.K., V.I.K., S.A., C.C., A.K., M.Ma., M.Mä. and N.D.W.; The manuscript was written by M.G., L.v.D., C.J.H. and L.S. with contributions from N.D.W., J.M.D., S.I., M.Ma., R.A., T.J., V.N.K., V.I.K., S.A., C.C., A.K. and M.Ma.; The manuscript was edited by M.G., C.J.H., L.v.D., L.S., K.T., T.K., M.H.D.L., C.L.S., S.I., A.K., S.A., N.D.W., M.Mä., C.C., H.K., B.B., O.B., S.S., T.S., A.S., J.M.D., L.P. and J.E.R.. All authors read and approved the final manuscript.

## Availability of data and materials

Sequencing data will be available upon publication at the European Nucleotide Archive (ENA) under the accession number PRJEB76829.

## METHODS

## 1. Laboratory work

### 1.1. Sample preparation and DNA extraction

All samples stem from skeletal human remains recovered from archaeological sites situated in Europe. Samples were not selected a priori for screening for *Human betaherpesvirus 6* species but are part of larger human population genetics projects and untargeted screening efforts to detect microbial pathogens in historical populations. As such most of the samples are teeth, with the exception of one incus bone, two petrous bones and a maxilla fragment (see Fig. S4).

Apical tooth roots were cut off with a drill and used for extraction since root cementum has been shown to contain more endogenous DNA than crown dentine ^50^. The root pieces were used whole to avoid heat damage during powdering with a drill and to reduce the risk of cross-contamination between samples. Contaminants were removed from the surface of tooth roots by soaking in 6% bleach for 5 minutes, then rinsing three times with milli-Q water (Millipore) and lastly soaking in 70% ethanol for 2 minutes, shaking the tubes during each round to dislodge particles. Finally, the samples were left to dry under a UV light for 2 hours. Petrous bones were sampled by drilling or cutting a sample from the core and if inner ear bones were present, these were used insteads. Petrous core and inner ear bones were decontaminated the same way as tooth roots.

Next, the samples were weighed, [20 * sample mass (mg)] µl of EDTA and [sample mass (mg) / 2] µl of proteinase K was added and the samples were left to digest for 72 hours on a rotating mixer at room temperature to compensate for the smaller surface area of the whole root compared to powder.

The DNA solution was concentrated to 250 µl (Vivaspin Turbo 15, 30,000 MWCO PES, Sartorius) and purified in large volume columns (High Pure Viral Nucleic Acid Large Volume Kit, Roche) using 2.5 ml of PB buffer, 1 ml of PE buffer and 100 μl of EB buffer (MinElute PCR Purification Kit, QIAGEN).

### 1.2. Library preparation

Double-stranded libraries were built using a protocol modified from the manufacturer’s instructions of the NEBNext® End Repair Module (E6050L, NEBNext), NEBNext® Quick Ligation Module (E6056L, NEBNext), and Illumina-specific adaptors ^51^, following established protocols ^51–53^. The end repair module was implemented using 30 μl of DNA extract, 12.5 μl of water, 5 μl of buffer and 2.5 μl of enzyme mix, incubating at 20 °C for 30 minutes. JDS067 and EDI001 (tooth sample) were processed in the ancient DNA lab of the Department of Archaeology, University of Cambridge and instead of 30 µl, 50µl were input for library preparation. The samples were purified using 500 μl PB and 650 μl of PE buffer and eluted in 30 μl EB buffer (MinElute PCR Purification Kit, QIAGEN). The adaptor ligation module was implemented using 10 μl of buffer, 5 μl of T4 ligase and 5 μl of adaptor mix ^51^, incubating at 20 °C for 15 minutes. The samples were purified as in the previous step and eluted in 30 μl of EB buffer (MinElute PCR Purification Kit, QIAGEN). The adaptor fill-in module was implemented using 12,2 μl of water, 5 μl of buffer, 0.8 μl dNTPs (25 mM each; Thermo Scientific) and 2 μl of Bst DNA polymerase, incubating at 37 °C for 30 and at 80 °C for 20 minutes. The libraries were amplified and combinatorial indexes (NEBNext Multiplex Oligos for Illumina, New England Biolabs) were added by PCR using HGS Diamond Taq DNA polymerase (Eurogentec). The samples were purified and eluted in 35 μl of EB buffer (MinElute PCR Purification Kit, QIAGEN). Three verification steps were implemented to make sure library preparation was successful and to measure the concentration of dsDNA/sequencing libraries – fluorometric quantitation (Qubit, Thermo Fisher Scientific), parallel capillary electrophoresis (Fragment Analyser, Agilent Technologies) or automated electrophoresis (Agilent TapeStation system, Agilent Technologies) and qPCR (KAPA Library Quantification Kit - Illumina® platforms).

### 1.3. Target Enrichment

The libraries were enriched using a Daicel Arbor Biosciences Custom myBaits multispecies viral capture kit, which contained, amongst others, sequences for all human herpesviruses, including 193 HHV-6 sequences and four HHV-7 sequences. Nine libraries were reamplified using KAPA Hifi Hotstart ReadyMix (2X) and primers IS5 and IS6 ^51^ prior to enrichment (SG1R input libraries in Table S4). Samples EDI001, EZH008, JDS067 and ORD001 were enriched using the v4 myBaits reagent kit and samples VAL004, KRJ003, KUU029, OLV143, OLV225, OLV303, OLV370 with the v5 myBaits reagent kit. Samples were each enriched in a single reaction with 2.75µl of baits per reaction. Enriched libraries were amplified using KAPA HiFi HotStart ReadyMix (2X) DNA Polymerase and primers IS5 and IS6 ^51^. For samples ORD001 and JDS067, previously enriched libraries were used for a second round of capture (see Table S4), due to low initial yield. Following amplification, the libraries were sequenced on an Illumina NextSeq500 sequencing platform (MID150, PE) at the University of Tartu, Institute of Genomics Core Facility, with other capture samples.

## 2. Human betaherpesvirus 6 Genomic Analysis

### 2.1. Ancient Datasets: Raw FASTQ preparation

Raw sequencing data (see Table S4) were merged by library and type of data (shotgun/capture). Data quality was assessed with fastqc ^54^. For paired-end data cutadapt ^55^ was run in paired-end mode with the pair-filter on “any”, quality trimming was set to 20 and reads under 20bp were discarded (--times 3 -e 0.2 -j 0 --trim-n). Reads were merged using Flash2.0 ^56^. Single-ended datasets were also trimmed and quality filtered using cutadapt ( -m 20 --nextseq-trim=20 --times 3 -e 0.2 -j 0 --trim-n). Dataset quality was assessed with multiqc^57^.

### 2.2. Ancient Datasets: Metagenomic Screening

Following data filtering, trimming and, if needed, paired-end read merging, samples were analysed using the taxonomic classifier KrakenUniq ^26^ to detect the presence of human associated pathogens. The database used for the analysis was composed of the following: dusted complete genomes and chromosome-level assemblies of bacteria, viruses, archaea, and protozoa, the human genome, the NCBI Viral Neighbor database, and the contaminant databases UniVec and EmVec. Heatmaps were computed using plotly ^58^, pandas, matplotlib^59^and numpy ^60^. E-values were calculated as follows 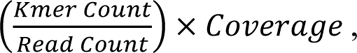 where “Coverage” is the coverage across the taxon kmer dictionary. The E-value cutoff to choose taxa for further inspection was 0.001. Taxa identifications were then verified by mapping the data to appropriate reference sequences as detailed for our final alignments, with the exception that BAM files were realigned using Picard ^61^.

### 2.3. Modern Datasets: Data preparation and metagenomic screening

Modern datasets were downloaded from NCBI and fragmented using the script FASTA-FRAG ^62^ into 50 bp fragments tiled across the sequences 20x with two base pair gaps. Fragments were analysed with kraken2 ^63^ to verify the species associated with the samples.

### 2.4. Comparative Mappings

Following the identification of the pathogens, target enrichment and further sequencing, datasets were merged by library and separately mapped to the HHV6A (NC_001664.4), HHV-6B (NC_000898.1) and HHV-7 (NC_001716.2) reference genomes (see Fig. S10-S13) using bwa aln (-n 0.04 -l 1000) ^64^ with samse for single-end data and merged paired-end reads. SAM files were converted to BAM format, sorted, indexed and filtered for mapped reads using samtools ^65^. Picard ^61^ was used to remove duplicates with the MarkDuplicates module. Following duplicate removal, datasets were merged by sample. Misincorporation patterns were computed and recalibration was done using mapDamage (v2.2.1) ^66^. Mapping plots were visualised using ^67^ (see Fig. 2), and mapping statistics were computed using Qualimap ^68^ and pysam ^69^, pandas, biopython ^70^ and numpy ^60^.

### 2.5. Phylogenetic analysis

Following mapping, consensus sequences were generated using freebayes (v.1.3.5) ^27^ and bcftools (v1.16 using htslib v1.6) ^71^ with either the reference sequence for HHV-6A or HHV-6B based on the species identification for each sample. Using deduplicated and rescaled BAM files for ancient DNA samples, SNPs were called with the following options: --max-complex-gap -1 --report-monomorphic -q 30 --min-alternate-count 3 --min-coverage 3 -m 30 -F 0.9 --ploidy 1. For modern samples, deduplicated BAM files were used with the following options: --max-complex-gap -1 --report-monomorphic --min-alternate-count 5 --min-coverage 5 -m 30 -F 0.9 --ploidy 1. VCF files were then filtered with bcftools filter with the options “-s LOWQUAL -i ’(QUAL>=30 && INFO/DP>=3) | (FORMAT/QR>100 && INFO/DP>=3) ’” for ancient DNA samples and “-s LOWQUAL -i ’(INFO/DP>=10)’” for modern DNA samples. Indels were removed. Filtered VCF files were then compressed using bcftools view and indexed using bcftools index.

Using bedtools intersect ^72^, SNPs which were only reported for aDNA samples were identified and visually inspected using IGV ^73^. This was also done for C>T and G>A variants to exclude possible ancient DNA misincorporations to be included in the final SNP alignment. Consensus sequences were called using bcftools consensus “-a “N” --exclude ’FILTER=“LOWQUAL”’” for modern samples and “-a “N” --exclude ’FILTER=“LOWQUAL” | INFO/DP<3’” for ancient DNA samples. Masking was applied for both modern and ancient DNA sequences to exclude repetitive intervals (see SI). Initial masking for the reference sequences for HHV-6A (NC_001664.4) and HHV-6B (NC_000898.1) was done based on the annotation provided by NCBI, by masking repeats. Additionally, these intervals were corrected by running GenMap on both reference sequences to adjust the interval coordinates. Intervals with low mappability preceding and following known repetitive regions were also included in the mask (see SI).

Masked consensus sequences and the reference sequence for each species were concatenated based on the identified taxon. Recombination analysis was performed in RDP5 (v4.45) ^41^. Initially, alignments were checked with a quick Phi test, which revealed “very good evidence for recombination” in both alignments. Recombination intervals were identified (see SI) and masked in the alignment. Using Mega11 ^74^, the alignment was filtered for variant positions present in 95% of all sequences. This alignment was then used to generate a maximum likelihood phylogeny in IQ-Tree2 ^42^ using the integrated “Model Finder Plus” without restrictions to choose the best-fitting models. Branch support was calculated with 10.000 ultrafast bootstrap approximation and SH-like approximate likelihood ratio test replicates.

### 2.6. Consensus Sequences Building for Temporal Analysis

For our temporal analysis, we mapped each ancient genome to a representative genome of the clade it clustered in, based on our maximum likelihood phylogeny (Fig. 3 and 4). All strains, with the exception of OLV225, clustered in an integrated clade, which evolved in a clonal manner, hence with an expectation of no larger structural changes. Consistently, recombination analysis of only integrated clades also revealed no evidence for recombination (see SI). Using the same approach as described above, consensus sequences were generated based on the following reference sequences: MT508932 (Clade A2), MT508930 (Clade A3), MT508933 (Clade A4), MT508941 (Clade B5) and MT508915 (Clade B8). OLV225 was mapped to its phylogenetically closest genome, based on our phylogeny, KY315534 (HP40E6). For a detailed description of the temporal analysis please see the supplementary information.

## 3. Human Data Genomic Analysis

### 3.1. Mapping

Before mapping, the sequences of adaptors and indexes and poly-G tales occurring due to the specifics of the NextSeq 500 technology were cut from the ends of DNA sequences using cutadapt 2.1 ^55^. Sequences shorter than 30 bp were also removed with the same program to avoid random mapping of sequences from other species. For paired-end sequencing data, R1 and R2 reads were merged using flash 1.2.11 ^56^. The sequences were mapped to reference sequence GRCh37 (hs37d5) using Burrows-Wheeler Aligner (BWA 0.7.17) ^64^ and command aln with re-seeding disabled. After mapping, the sequences were converted to BAM format and only sequences that mapped to the human genome were kept with samtools 1.9 ^65^. Next, data from different flow cell lanes was merged and duplicates were removed with picard 2.20.8 (http://broadinstitute.github.io/picard/index.html). Indels were realigned with GATK 3.5 ^75^ and lastly, reads with mapping quality under 10 were filtered out with samtools 1.3 ^65^.

### 3.2. aDNA authentication

As a result of degradation over time, aDNA can be distinguished from modern DNA by certain characteristics: short fragments and a high frequency of C=>T substitutions at the 5’ ends of sequences due to cytosine deamination. The program mapDamage2.0 ^66^ was used to estimate the frequency of 5’ C=>T transitions. mtDNA contamination was estimated using the method from Fu et al. 2013 ^76^. This included calling an mtDNA consensus sequence based on reads with mapping quality at least 30 and positions with at least 5x coverage, aligning the consensus with 311 other human mtDNA sequences from Fu et al. 2013 ^76^, mapping the original mtDNA reads to the consensus sequence and running contamMix 1.0-10 with the reads mapping to the consensus and the 312 aligned mtDNA sequences while trimming 7 bases from the ends of reads with the option trimBases. For the male individuals, contamination was also estimated based on chromosome X using the two contamination estimation methods first described in Rasmussen et al. 2011 ^77^ and incorporated in the ANGSD software ^78^ in the script contamination.R.

### 3.3. Calculating general statistics and determining genetic sex

Samtools 1.3 ^65^ option stats was used to determine the number of final reads, average read length, average coverage etc. Genetic sex was calculated using the script sexing.py from Skoglund et al. 2013 ^79^, estimating the fraction of reads mapping to Y chromosome out of all reads mapping to either X or Y chromosome.

