## Supplementary Information for "2500 Years of Human Betaherpesvirus 6A and 6B Evolution Revealed by Ancient DNA"

**I. Archaeological Data** 2
a) Sint-Truiden, Limburg, Belgium (by Natasja De Winter) 2
b) Hospital of St. John, Cambridgeshire, United Kingdom (by Jenna Dittmar, Sarah Inskip and Craig Cessford) 2
c) Edix Hill, Barrington A, Cambridgeshire, United Kingdom (by Jenna Dittmar, Sarah Inskip and Craig Cessford) 3
d) Kukruse, Ida-Viru County, Estonia (by Martin Malve, Tõnno Jonuks) 4
e) Karja, Saaremaa, Estonia (by Marika Mägi, Martin Malve and Raili Allmäe) 5
f) Valjala, Saaremaa, Estonia (by Marika Mägi, Martin Malve and Raili Allmäe) 5
g) Ezhol, Kortkerossky district, Komi Republic, Russia (by V. N. Karmanov, V. I. Khartanovich and Aivar Kriiska) 6
h) Ordona, Apulia, Italy 6
**II. Radiocarbon Dating** 7
**III. Masking Intervals For Human Betaherpesvirus Reference Sequences** 7
**IV. Human Betaherpesvirus Recombination Analysis** 7
**V. Human Betaherpesvirus SNPEff Analysis** 7
**VI. Human Genome Clinvar Analysis (by Lehti Saag)** 8
**VII. Human Betaherpesvirus Temporal Analysis (by Lucy van Dorp)** 8
**VIII. SUPPLEMENTARY FIGURES** 11
**IX. SUPPLEMENTARY INFORMATION REFERENCES** 28

### I. Archaeological Data

#### a) Sint-Truiden, Limburg, Belgium (*by Natasja De Winter*)

(OLV143, OLV225, OLV303, OLV370)

From 2018 to 2020, "Aron bv" conducted an excavation in the city centre of Sint-Truiden [1,2], following the redevelopment of the Trudoplein, the Groenmarkt and the streets surrounding it. In the northern part of the excavated area, the Trudoplein borders the tower of the former Sint-Trudo abbey to the south. The Groenmarkt in turn borders the Trudoplein to the south. East of this market is the parish church, the Church of Our Lady, which dates back to the middle of the 11th century. To the south, the market is bordered by the town hall.

Archaeological evidence dating back to the Early Middle Ages were identified at both Trudoplein and the Groenmarkt. These sites are probably linked to the settlement that developed around the monastery and was founded by Trudo, a nobleman of Frankish descent known for his piety and missionary work, in the second half of the 7th century on the family estate of *Sarchinium*. The existence of such a settlement was already mentioned by Donatus in his *Vita Sancti Trudonis* in the second half of the 8th century. Thus, the excavation seems to confirm this.

Soon after, the investigated area was used for burials and this remained unchanged for centuries. Consequently, the vast majority of the archaeological traces found within the site consist of graves. In those burials, a total of 3,046 different individuals were identified, both male and female. Based on radiocarbon dating, the oldest individuals date back to the late 7th century or the 8th century. In the early Middle Ages, the burial field extended over a large area: from the abbey tower to today's town hall, a distance of approximately 100 metres. It must have been a part of the burial ground belonging to the abbey since there are no other churches known from this period. On today's Trudoplein, burials continued to take place around the abbey tower. The youngest grave in the area near the tower dates to the 14th century; no burials were made here afterwards. On the Groenmarkt burials also continued after the year 1000, but the cemetery situated there moved further eastward. In the second half of the 11th century, the Church of Our Lady was built east of the Groenmarkt. From the end of the 13th century, possibly a little earlier, burials were only carried out within the cemetery wall surrounding this church, a wall that bisected the Groenmarkt from north to south. The western part of the Groenmarkt must have been used as a market from then on. The eastern part included a section of the cemetery around the Church of Our Lady, which was finally given up in the second half of the 18th century.

OLV143 (SK0132) (Fig. S1), OLV225 (SK1197) (Fig. S2) and OLV303 (SK1917) (Fig. S3) are all female adults, excavated at the Trudoplein and the Groenmarkt (Zones 1 and 3). Whereas OLV370 (SK2326) (Fig. S4) is the only male adult for which *Human betaherpesvirus 6* sequences could be identified, the skeleton was excavated at the Groenmarkt (Zone 2) and was recovered with an unusual right arm position, indicating a pit deposition.

#### b) Hospital of St. John, Cambridgeshire, United Kingdom

(*by Jenna Dittmar, Sarah Inskip and Craig Cessford*)

(JDS067)

The Hospital of St. John the Evangelist, Cambridge, United Kingdom, was founded in the town of Cambridge at the very end of the 12th century by the townsfolk, with burial rights acquired in the early 13th century, and was in use until the early 16th century when it was dissolved to create St John's College. The Hospital was established for the care of the poor and infirm, and has been well studied using ancient DNA [3,4]. Over 400 complete or partial articulated skeletons were recovered by the Cambridge Archaeological Unit in 2010/2011 [5]. Subsequent radiocarbon dating has broadly confirmed and refined the dating of the cemetery.

JDS067 (PSN192), who was genetically sexed as male, was a boy between the ages of 8–10 years of age at the time of his death, based on the development of his dentition and skeleton. He lived during the 14th century and was buried in the cemetery. A single *Herpes simplex 1 virus* genome (JDS005) has been published from the site in 2022 [6]. Between 50–75% of the JDS067 (PSN192) skeleton was recovered during excavation, with the skeletal elements of the lower limbs missing. This individual had active cribra orbitalia at the time of his death. The etiology of cribra orbitalia remains under debate but most research suggests these lesions are associated with one of the anemias [7], though some research suggests that these lesions can also be caused by other conditions including B12 deficiency [8,9]. Other indicators of physiological stress included evidence of linear enamel hypoplasia, permanent defects in dental enamel that result from a disruption during amelogenesis [10,11]. This indicates that this child experienced at least one episode of arrested growth disruption between the ages of 1 and 6 years of age. Numerous aetiologies for LEH have been identified (see [12]) including: vitamin A and D deficiencies [13,14], infections, malnutrition [15], low birth weight [16–18], as well as a variety of specific conditions and disturbances [19–21]. The bodies of the thoracic and lumbar vertebrae had reduced cortical bone density, resulting in the vertebral bodies having a 'lace-like' appearance. Cortical defects were also present on the superior facets of the axis (C2) which corresponded with similar defects on the inferior facets of the atlas (C1). This individual has no evidence for vitamin C or D deficiency.

#### c) Edix Hill, Barrington A, Cambridgeshire, United Kingdom

(*by Jenna Dittmar, Sarah Inskip and Craig Cessford*)
(EDI001)

The Anglo-Saxon cemetery of Edix Hill, located between the Cambridgeshire villages of Barrington and Orwell was initially discovered in the 19th century. Excavations between 1989 and 1991 revealed part of an inhumation cemetery comprising 149 individuals in 115 graves, dating to between 500 and 625/650, although one burial was believed to be possibly Iron Age [22]. It is estimated that there may originally have been around 300 burials in total with a complete cross-section of the population by age and sex suggesting that the burials relate to a community of 50–65 people spanning around 150 years. The initial dating of the cemetery was based primarily upon artefact typologies of the various grave goods and seriation by correspondence analysis, with burials broadly divided into earlier and later groups and some evidence for spatial patterning over time. Subsequent radiocarbon dating has broadly confirmed and refined the dating of the cemetery.

EDI001 (PSN554/Sk405/Grave 76), who was genetically sexed as male, was between the age of 16–18 years of age at the time of his death based on the development of his dentition and skeleton. The skeleton was accompanied by a buckle and a spearhead (see Fig. 3.79 in [22]). This individual dates to the 6th–7th century, with the identification of the plague bacterium *Yersinia Pestis* indicating a mid-6th-century date[23]. In similarity to the Hospital of St. John, the Anglo-Saxon site of Edix Hill has also been well studied via ancient DNA. It is particularly well known as a First Plague Pandemic site, with numerous plague genomes [23] and a *Haemophilus influenzae serotype b*/*Yerinia pestis* co-infection (EDI064) [24] having been identified on site. Individual EDI001 (PSN554) has evidence that they experienced at least one period of growth disruption (as indicated by the presence of LEH) during their early childhood, but neither cribra orbitalia, nor porotic hyperostosis were observed. There was, however, new subperiosteal bone formation located inside the nasal aperture. Mixed (both active and healed) subperiosteal bone formation was present on the posterior aspect of the right tibia and fibula. Due to taphonomic damage, it was not possible to assess the left lower limb skeletal elements. New subperiosteal bone formation was present on the shaft of the left second metatarsal. Subperiosteal new bone formation is caused by inflammation of the periosteum (the connective tissue that covers the bone surface, except for the joint surfaces) that can be initiated by infectious diseases or trauma. An area of bone destruction was identified between the superior talar facet of the left calcaneus and corresponding calcaneal facet on the talus. This individual's femur also had a possible bending deformity, which would indicate that this individual experienced a period of chronic vitamin D deficiency during his childhood. Similarly to the previously reported coinfection detected for individual EDI064, individual EDI001 was also infected by the plague bacterium *Yersinia pestis* [23], but the HHV-6 carriage described in this study was not reported at the time.

#### d) Kukruse, Ida-Viru County, Estonia (*by Martin Malve, Tõnno Jonuks*)

(KUU029)

Kukruse cemetery is a burial ground located in Northeast Estonia [25]. The earliest stratum of the site is formed by a cremation cemetery from the 1st millennium CE, followed by an inhumation cemetery right at the same place. The inhumation cemetery was used mostly during the 12th–13th century CE, as attested by the grave goods, such as pottery, decorative pins, knife sheaths and coin pendants; however, single inhumations were done as late as the 17th century. The individual studied here is dated between 1290 and 1400 cal CE with 95.4% probability (620±30; Beta-293533; Lõhmus et al. 2011; calibrated with OxCal v4.4.4.).

A part of the cemetery with a total of 40 inhumation graves, including men, women and children, was rescue excavated in 2009 and 2010. Some of the individuals were buried in wooden coffins and in some cases, above-ground markings of the graves were documented. Some of the deceased – especially a group of five inhumations in the eastern part of the cemetery – were interred with rich funerary equipment, ranging from weapons and tools to personal attires and jewellery. Others were accompanied by personal items and dress accessories, and some individuals were buried with no grave goods at all. In many cases ceramic vessels, that initially also contained food, had been placed at the head or foot of the graves.

The majority of the graves at the Kukruse cemetery belong to the transition period in Estonia. During the late 12th century, the first Crusades extended to this part of Estonia and the region was conquered and officially Christianised in the 1220s. Part of this process was also the emergence of new inhumation cemeteries, where most of the population of a community was buried. Differently from the previous periods, we know a number of non-adult inhumations since the late 12th/early 13th century. In total, the non-adult population at the Kukruse cemetery is 41%, while infants up to 2 years of age form 1/5th of the cemetery population [26].

The grave was located at the western part of the cemetery and had been dug through the cremation cemetery stratum. It is directly dated to 1290 to 1400 cal BCE (620±30; Beta-293533) [25], demonstrating that this burial was conducted after the official Christianization of the region (this also is in accordance with absent grave goods and body's orientation in the grave). Wood fragments – possibly a log that once marked the grave – were found exactly above the inhumation. A fully preserved and articulated skeleton of a non-adult individual was discovered from the grave. It was embedded on its back with both upper and lower limbs extended, and its head directed towards the west. No grave goods accompanied the deceased. The age of death was estimated to be between 10.5

months to 1.5 years based on the development and eruption of the teeth [26]. The individual is genetically sexed as a female.

#### e) Karja, Saaremaa, Estonia (*by Marika Mägi, Martin Malve and Raili Allmäe*) (KRJ003)

Archaeologist Aita Kustin excavated the Karja cemetery on Saaremaa island, Estonia, between the buildings belonging to the later Karja manor complex in 1955 [27]. She unearthed 34 skeletons, but the actual number of burials would have been around 70. The excavated skeletons belonged mainly to the period between 1200–1300 and were buried with their heads to the West, in a Christian manner. An empty area between two groups of graves probably marks a no longer visible wooden church or chapel. Several graves contained artefacts of local style, mainly jewellery and other attributes. The deceased were probably inhabitants and household members of the Medieval Karja manor. Written sources, written some centuries later, indicate the Karja manor played a central role in the district.

KRJ003 (Burial No 3 –1955) [28] was male, died at the age of 30–35 years. He had been buried between 1220 and 1260 AD, in a coffin, with head to West-South-West, in a 40–50 cm deep grave. An iron knife was found on his right hip bone. The male individual had a calculated body height of 172.4 cm and a weight of 73.3 kg. The muscle attachments on the skeleton, especially on the bones of his upper arms, are well developed. Rotator cuff disease can be observed on both shoulder joints but is more pronounced on the right side. Osteoarthritis of the elbow joint is also noticeable on the right side. The observed muscle attachment patterns, in combination with pathological changes and osteological measurements of upper limb bones, likely reflect repeated and stronger use of the right arm. Further, we observed a healed fracture of the distal part of the right radius. The skeleton is stored at Tallinn University, Archaeological Research Collections, AI 4115.

#### f) Valjala, Saaremaa, Estonia (*by Marika Mägi, Martin Malve and Raili Allmäe*) (VAL004)

Excavation at Valjala took place in 2010 [29], when 26 skeletons were unearthed 45–50 m from the Medieval stone church at Valjala, Saaremaa island, the largest Island on the Estonian Baltic Sea coast. Burials belonged mainly to the period 1200–1300 CE and may have been originally situated in the churchyard of a since-disappeared wooden church or chapel. Another cluster of similar, mainly 13th-century burials, have been uncovered around the earliest stone church. All burials excavated in 2010 were presumably Christian, with heads oriented to the West or North-West. Several graves, especially those of women, contained artefacts that belonged to local styles. Some female burials included numerous ornaments and metal attributes, indicating the wealth and status of the deceased. This ostensible wealth is supported by available 13th-century chronicles, where Valjala was called the very centre of the Saaremaa island.

VAL004 (Burial No 4 – 2010; Valjala IV) at Valjala belonged to a female (see Fig. S3), who died at 40–50 years of age, buried with her head to the West, in the late 12th century CE. Only the upper part of the burial, down to the pelvis, was excavated; the rest had been destroyed by road constructions prior to the excavation. She wore a head-dress decorated with elaborate metal ornamentation, a chain arrangement fixed with two dress pins, a penannular brooch and other ornaments. Additional ornaments had been placed on the left hip of the deceased. The female individual had been buried in a wooden coffin, lined and covered with stones. Her grave is one of the earliest burials in the Valjala churchyard. During her lifetime the woman had suffered from diseases that accompany the process of ageing (e.g. osteoarthrosis). The skeleton is stored at Tallinn University, Archaeological Research Collections, AI 7585.

#### g) Ezhol, Kortkerossky district, Komi Republic, Russia

(*by V. N. Karmanov, V. I. Khartanovich and Aivar Kriiska*)
(EZH008)

The burial site of Ezhol is located in the Kortkerossky district, Komi Republic, Russia, on a sandy aeolian dune deposited on a river terrace bordering the old floodplain of the Vychegda River. The burial site was discovered in 2013 and partially excavated by V. N. Karmanov, and later – in 2014–2015 by T. Yu (total area of the excavations 422 m²). Eight round or oval burial mounds of up to 0.6 m in height and with dimensions of 4.0×3.1 m to 7.3×5.5 m, and two flat graves have been investigated. There were 1 to 3 burials beneath each of the mounds. A total of 18 burials were excavated – 16 under the burial mounds and two in flat graves [30]. The dead were placed in graves 1.5–2.8 m long and 0.4–1.75 m wide, lying flat on their backs with heads facing the river. Iron knives, boot decorations, some jewellery such as brooches and amber beads, as well as broken clay vessels were found in the graves. The burial site is dated to the second half of the 5th century and the first half of the 6th century CE [31]. Similar to other analogous burial sites with a combination of mounds and flat graves known in the basin of the Vychegda River and in the valley of the Izhma River, the Ezhol burial sites likely also belonged to nomadic hunter-fisher-gatherer groups which migrated from the Kama River basin [31]. The skeletons are stored in the Institute of Language, Literature and History, Komi Science Centre, Ural Branch, Russian Academy of Sciences.

Ancient DNA analysis for individual EZH008 was performed using a petrous bone of a 50-year-old woman from grave 2 of burial mound IV, which was found in 2015 [30]. In addition to the woman's bones, an iron knife and a bead made of a copper-containing alloy were also found in the grave. Next to grave 2 was the burial of a 20-30-year-old man (grave 1). According to the stratigraphy, grave 1 was created later than grave 2 and was dug in such a way as not to damage it; later, a single mound was made over both graves.

Radiocarbon (AMS) dating from EZH008 yielded an age of 1896±25 BP (UBA-45348), 76–218 calCE with a probability of 95.4% (calibration, [32,33]), but stable isotopes indicate that the date is affected by the freshwater reservoir effect. The nitrogen and carbon values ($\delta^{13}$C – -20.9‰, $\delta^{15}$N – 11.6‰) are particularly similar in terms of nitrogen of other analysed human bones from the Ezhol burial site, according to which it was suggested that freshwater fish constituted at least 25–30% of peoples diet [31]. Since the reservoir shift is not clear, the burial should rather be dated not to the 1st–3rd centuries CE but to a range that considers both the radiocarbon dating and the youngest possible age of the cemetery – from the second half of the 1st century to the first half of the 6th century CE.

#### h) Ordona, Apulia, Italy

(ORD001)

The necropolis of Herdonia is situated in today's Ordona, Apulian Foggia, Italy. The settlement of Herdonia is expected to have started from Neolithic, and the site became an important centre of the Daunian culture around the 6th century CE. The necropolis had been studied over multiple archaeological campaigns from 1978–1981 [34]. The individuals recovered during the excavation have been studied osteologically [35] and using ancient DNA analysis [36]. Individual ORD001 was also included but yielded only low level of endogenous DNA (0.0396, XX, mitochondrial haplotype H5c).

#### II. Radiocarbon Dating

All partial and full genomes described in this study have been radiocarbon dated. However, analysis was not performed in the same laboratory. Instead, the analysis was spread across five laboratories, since they were performed during the course of different projects from multiple European research institutions. More details for each date can be found in the Supplementary Information Table S10.

#### III. Masking Intervals For *Human Betaherpesvirus* Reference Sequences

For our analysis, we masked repetitive and low-complexity intervals for both HHV-6A and HHV-6B. This was done by using the NCBI annotation and correcting intervals using mappability estimates as calculated by GenMap [37] (map -K 30 -E 2). The masking intervals were defined as follows:

| | | | |
|---|---|---|---|
| NC_001664.4 | 0 - 8089 | NC_000898.1 | 0 - 8911 |
| NC_001664.4 | 127549 - 128233 | NC_000898.1 | 9315 - 9522 |
| NC_001664.4 | 131077 - 131854 | NC_000898.1 | 129039 - 129681 |
| NC_001664.4 | 138050 - 140951 | NC_000898.1 | 133524 - 134076 |
| NC_001664.4 | 151234 - 159378 | NC_000898.1 | 140078 - 142691 |
| | | NC_000898.1 | 153017 - 162114 |

#### IV. *Human Betaherpesvirus* Recombination Analysis

Recombination analysis was performed in RDP5 (v4.45) [38]. Initially, alignments were checked with a quick Phi test, which revealed very good evidence for recombination in both alignments. For our alignments to the HHV-6A reference sequence, RDP was run with a secondary scan by Bootscan with zero permutations and a minimum p-value of 0.001. For RDP the “inner reference” option and a window size of 60 were used. A minimum of four tools were needed for a recombination event to be detected. The rest of the options were left on default. The same settings were used for our HHV-6B analysis, with the exception that one event was rejected as it spanned the entire sequence and was uncertain. Alignments were then exported in FASTA format masked for recombinant intervals.

For HHV-6B recombination signals were quite unstable across tools and settings tested, which was seemingly driven, amongst others, by the more basal branches, which are heavily populated by acquired strains. A recombination analysis of only ici clades revealed no evidence for recombination amongst the ici clades of either species.

#### V. *Human Betaherpesvirus* SNPEff Analysis

To explore the functional impact the detected SNPs could have had on the viruses, we ran SNPEff for all our new ancient genomes above a mean depth of coverage of 3X. For HHV-6A we only detected two variants with predicted HIGH IMPACT for JDS067. Both variants are predicted to cause stop gain mutations in genes U24 and U71. However, it should be noted that JDS067 is our lowest coverage full genome. For HHV-6B, variants with predicted HIGH IMPACT were detected for EDI001 (two variants), EZH008 (two variants) and OLV225 (three variants) (see Fig. S14a-c and Table S5 and S6). Variants falling within masked intervals were not considered. Variants 23,328 (T>G) and 103,758 (G>A) are predicted stop loss/splice variant in U12 and a start loss variant in U67 respectively, and occur in EDI001 and OLV225, which could be due to annotation issues. For sample EZH008, a frameshift causing a deletion in U91 and a frameshift causing an insertion in B9 were detected. Finally for OLV225, a stop gain variant at 145,618 in U95 was detected.

While inactivating mutations are known to occur for HHV-6 [39], it should be noted that the annotation for herpesviruses are known to be rather complex and not fully resolved[40,41]. HHV-6A and HHV-6B are no exceptions. This makes the validation of our results more difficult than for newly sequenced modern genomes as we are working with ancient DNA. References for each integrated clade were mostly unannotated, which did not allow us to identify potential non-synonymous mutations within clades.

#### VI. Human Genome Clinvar Analysis (*by Lehti Saag*)

The clinical variant (ClinVar) analysis resulted in 0 to 78 likely pathogenic or pathogenic high-quality genotyped variants with disease name information present per individual and 98 in total. However, out of these, only one variant has a clear relevance for predisposition to infection: rs901844850. This variant is relevant for MHC class II deficiency, also known as Bare Lymphocyte Syndrome type II (BLS2) [42] and is genotyped as heterozygous in one individual (EDI001), with 6 C (reference) and 2 T (alternative) alleles present. Even though CT and GA allele pairs also make up most of the ClinVar variant list, we cannot be sure that the 2 T alleles in our sample are not the result of cytosine deamination instead of being true alleles.

The ClinVar VCF file (https://ftp.ncbi.nlm.nih.gov/pub/clinvar/vcf_GRCh37/) was downloaded on 17/12/2023. The file was filtered for variant type (CLNVC) "single_nucleotide_variant" and clinical significance (CLNSIG) "Likely_pathogenic" or "Pathogenic", resulting in 118,655 variants, out of which 10,568 had two alternative alleles present and 1,402 had three. Genotypes for these variants were called using GATK 3.5 [43] HaplotypeCaller with --genotyping_mode GENOTYPE_GIVEN_ALLELES and --output_mode EMIT_VARIANTS_ONLY. Next, the resulting per sample VCF files were filtered for only genotypes including an alternative allele (-e GT 0/0), with a depth of coverage of at least 2 (-i INFO/DP>=2) and mapping quality of at least 30 (-i INFO/MQ>=30) that pass the GATK low-quality filter (-e FILTER="LowQual"). Finally, heterozygous variants for haploid loci (mitochondrial variants for all individuals, X and Y chromosome loci for males) and variants for which the disease name (CLNDN) was not provided were filtered out.

#### VII. *Human Betaherpesvirus* Temporal Analysis (*by Lucy van Dorp*)

A number of approaches were taken to assess the extent of temporality in the dataset including the six HHV-6A aDNA samples (all integrated) and the three HHV-6B aDNA samples (two integrated and one likely acquired). The first approach focused exclusively on strains identified as 'acquired' conducting independent analysis for 6A and 6B the second considered 'all strains' independently for 6A and 6B.

In each case a core SNP-calling approach was used, employing the snippy-core toolkit (https://github.com/tseemann/snippy) which applies FreeBayes to identify SNPs present in 99% of the included isolates, providing a .bed coordinate file to mask regions identified as repetitive or hyper-variable, as previously described. Regions of the core alignment putatively deriving from recombination were evaluated using 3Seq[44] to identify recombinant regions when considering all possible triplet pairs in the alignment. All identified positions were subsequently masked before maximum likelihood phylogenetic trees were constructed using IQ-Tree 2[45] specifying 1,000 boot-strap iterations and a GTR+G substitution model. Resulting trees were assessed for the extent of temporal signal using the roototip() function implemented in BactDating[46], following pre-specification of the initRoot() functionality. When known, point estimates for collection date were used, whereas in other instances a uniform prior was used for samples without collection date but for which there was reasonable confidence on the date range for which sampling had taken place. For ancient samples, the mean radiocarbon estimates, were used as an initial prior on the sample age. In all cases the extent of temporal signal was evaluated using linear regression followed by assessment of significance following 10,000 permutations of the sampling date to produce an empirical p-value.

The procedure was repeated using an alternate method to correct for incongruent sites deriving from recombination or alternative mechanisms. In particular, maximum parsimony trees were constructed using MPBoot[47] with 10,000 ultra-fast bootstraps from the clean masked core alignment (clean.full.aln) in each case and the resulting phylogeny (see Fig. S15-S17) and alignment used to identify and screen for homoplasies using HomoplasyFinder[48], to stringently remove those sites deriving from recombination, poor alignment quality or post-mortem damage. A summary of the number of SNPs considered and the resulting temporality following the procedure described is given in Supplementary Table A below.

| **Analysis** | **Core SNPs following masking/recombination pruning** | **Global r2 (initRoot)** | **Empirical p-value (initRoot)** |
|---|---|---|---|
| Acquired strains 6A (n=12) | 8,039 | 0.05 | 0.30 |
| Acquired strains 6B (n=72) | 6,054 | 0.13 | 0.0004 |
| All strains 6A (n=66) | 107,943 | 0.03 | 0.12 |
| All strains 6B (n=181) | 34,771 | -0.38 | 0.92 |

| **Analysis** | **Core SNPs following masking/homoplasy pruning** | **Global r2 (initRoot)** | **Empirical p-value (initRoot)** |
|---|---|---|---|
| Acquired strains 6A (n=12) | 7874<br>(165 homoplasies) | 0.05 | 0.2856 |
| Acquired strains 6B (n=72) | 5156<br>(1043 homoplasies) | 0.15 | 0.02 |
| All strains 6A (n=66) | 107018<br>(1632 homoplasies) | 0.07 | 0.05 |
| All strains 6B (n=181) | 33864<br>(911 homoplasies) | 0 | 0.43 |

***Supplementary Table A:*** (top) Core SNPs considered following masking and recombination pruning of alignments for subsets of data (either acquired strains only or jointly considering acquired and integrated data) considered for temporal analysis. (bottom) Core SNPs considered following masking and homoplasy pruning of alignments for subsets of data (either acquired strains only or jointly considering acquired and integrated data) considered for temporal analysis.

We followed a similar approach to evaluate temporality of integrated clades which included at least one ancient observation (Supplementary Table B). Clades of integrated HHV-6A and HHV-6B were identified as per the phylogenetic relationships and with reference to Aswad et al. [49]. We recover a temporal signal in a single integrated clade (A2) which is largely driven by the placement of JDS067 and was not stable following different recombination pruning methods. Hence a formal phylogenetic tip-dating approach was not implemented in this case.

It is likely that additional data, particularly from older data or supporting a within clade time-series may offer future opportunities to employ phylogenetic methods to calibrate the age of these clades.

| **Analysis** | **Core SNPs following masking/homoplasy pruning** | **Minimum age of clade based on ancient samples** | **Temporal r2 / p-value** |
|---|---|---|---|
| B5 integration (n=14) | 6,963<br>(28 homoplasies) | 1497 +- 24 BP | 0.31 / 0.076 |
| B8 integration (n=48) | 19,359<br>(55 homoplasies) | 1896 +- 25 BP | -0.01 / 0.812 |

| **Analysis** | **Core SNPs following masking/homoplasy pruning** | **Minimum age of clade based on ancient samples** | **Temporal r2 / p-value** |
|---|---|---|---|
| A2 integration (n=21) | 96,517<br>(25 homoplasies) | 669BP +- 24 BP | 0.99 / 0.004<br>Mu 1.57e-01<br>Root date -1297.98 |
| A3 integration (n=12) | 29,725<br>(28 homoplasies) | 1285 +- 24 BP | 0.01 / 0.146 |
| A4 integration (n=20) | 21,932<br>(116 homoplasies) | 1161 +- 23BP | -0.002 / 0.99 |

***Supplementary Table B:*** Core SNPs considered following masking and homoplasy pruning of alignments for subsets of data (by integrated clade) considered for temporal analysis.

#### VIII. SUPPLEMENTARY FIGURES

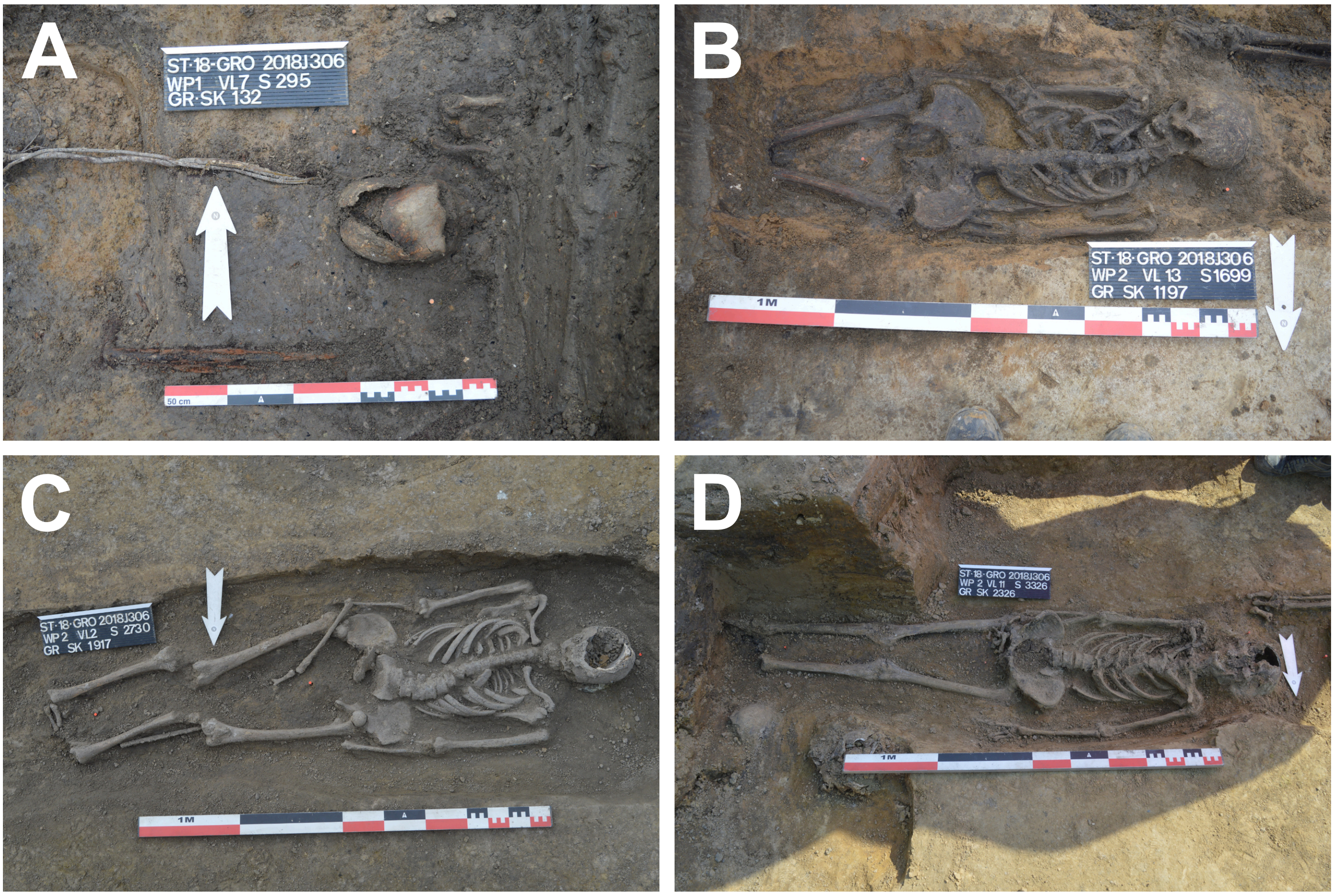


**Figure S1**: Images of the burials in situ for a) OLV143; b) OLV225; c) OLV303 and d) OLV370 from the Sint-Truiden excavations (Photographs by Natasja De Winter).

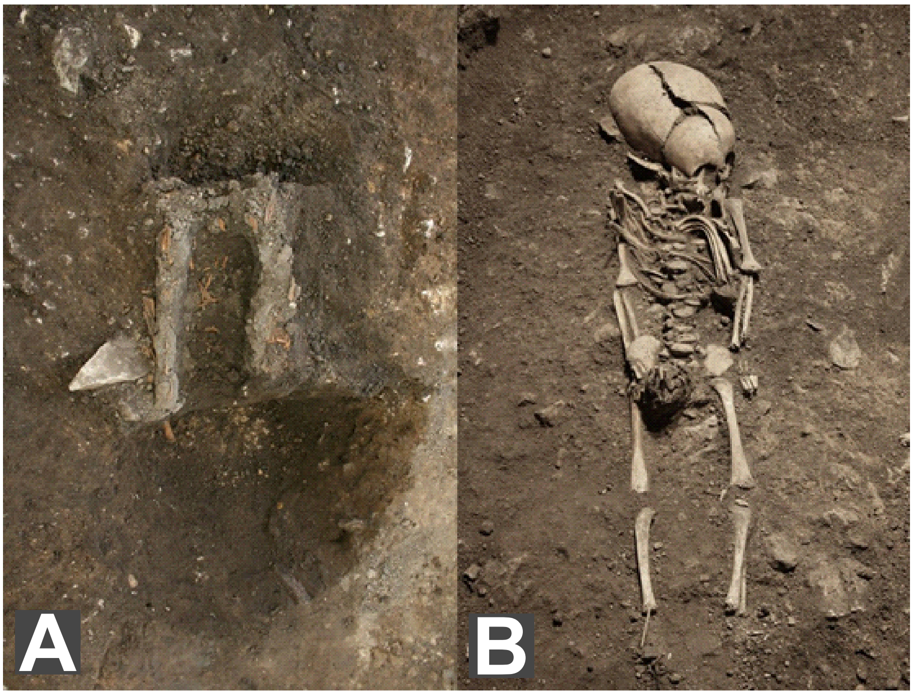


**Figure S2:** Grave of individual Kukruse XXXI (KUU029): a) wooden log fragments that were most probably used as above-ground grave marker and b) the skeletal remains of the infant (Photograph by Tõnno Jonuks).

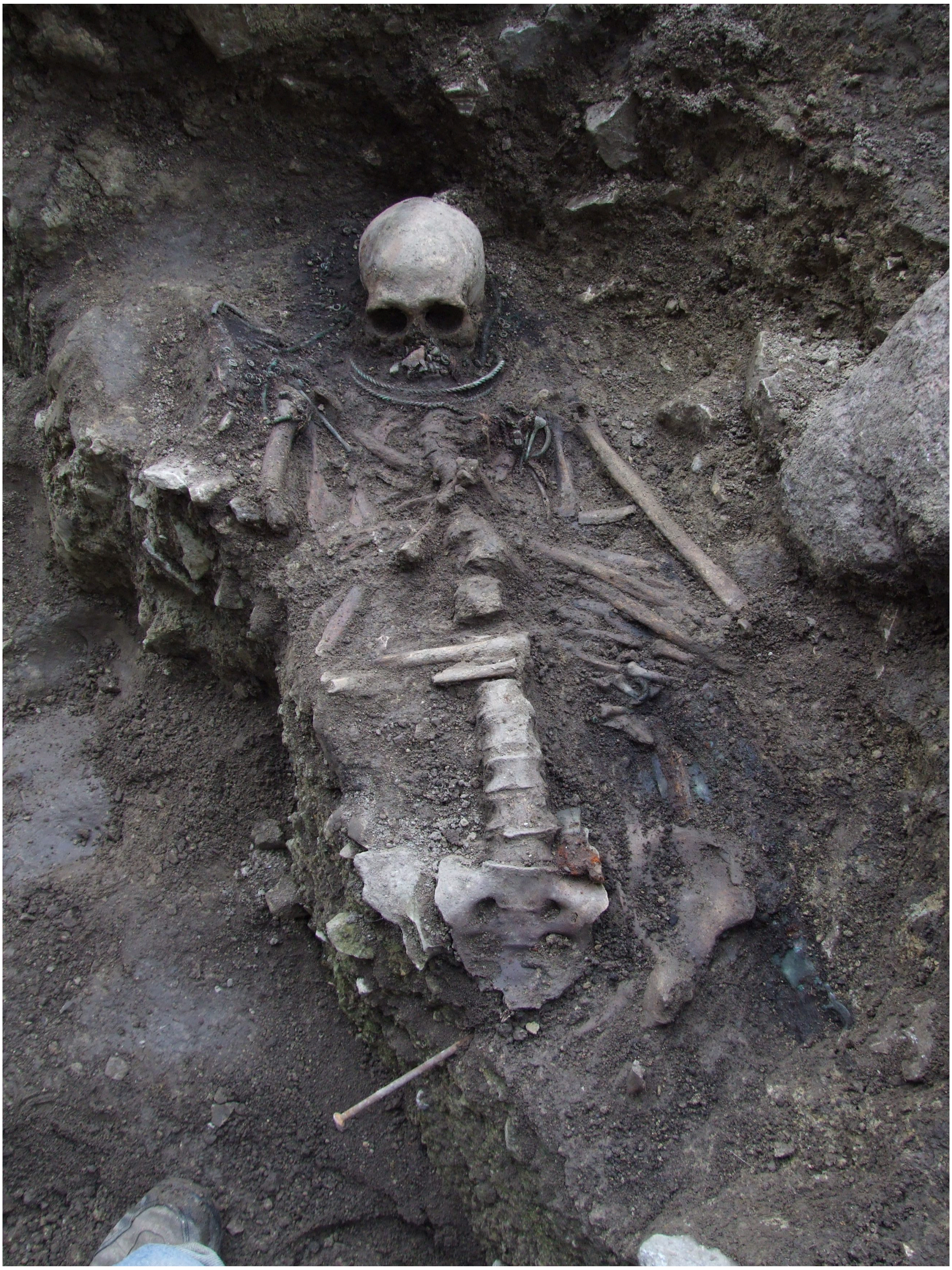

**Figure S3:** Grave No 4 (VAL004) during excavations at Valjala in 2010
(Photograph by Marika Mägi).

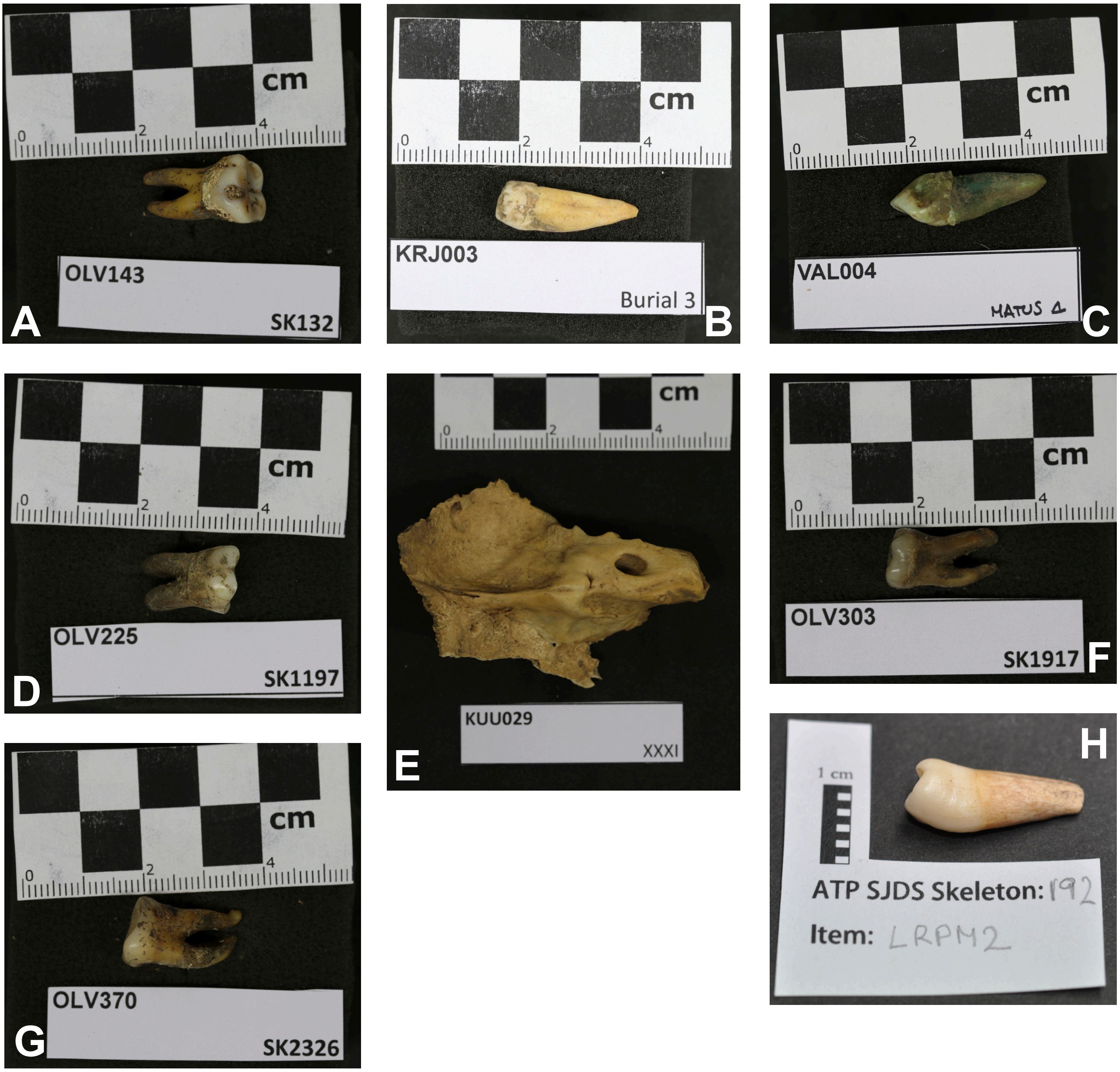


**Figure S4:**

Images taken of analysed samples upon entry in the ancient DNA laboratory at the University of Tartu and at the University of Cambridge. A) OLV143 (Photograph by Helja Kabral); B) KRJ003 (Photograph by Stefania Sasso); C) VAL004 (Photograph by Stefania Sasso); D) OLV225 (Photograph by Helja Kabral); E) KUU029 (Photograph by Stefania Sasso); F) OLV303 (Photograph by Helja Kabral); G) OLV370 (Photograph by Helja Kabral) and H) JDS067 (Photograph by Sarah A. Inskip).

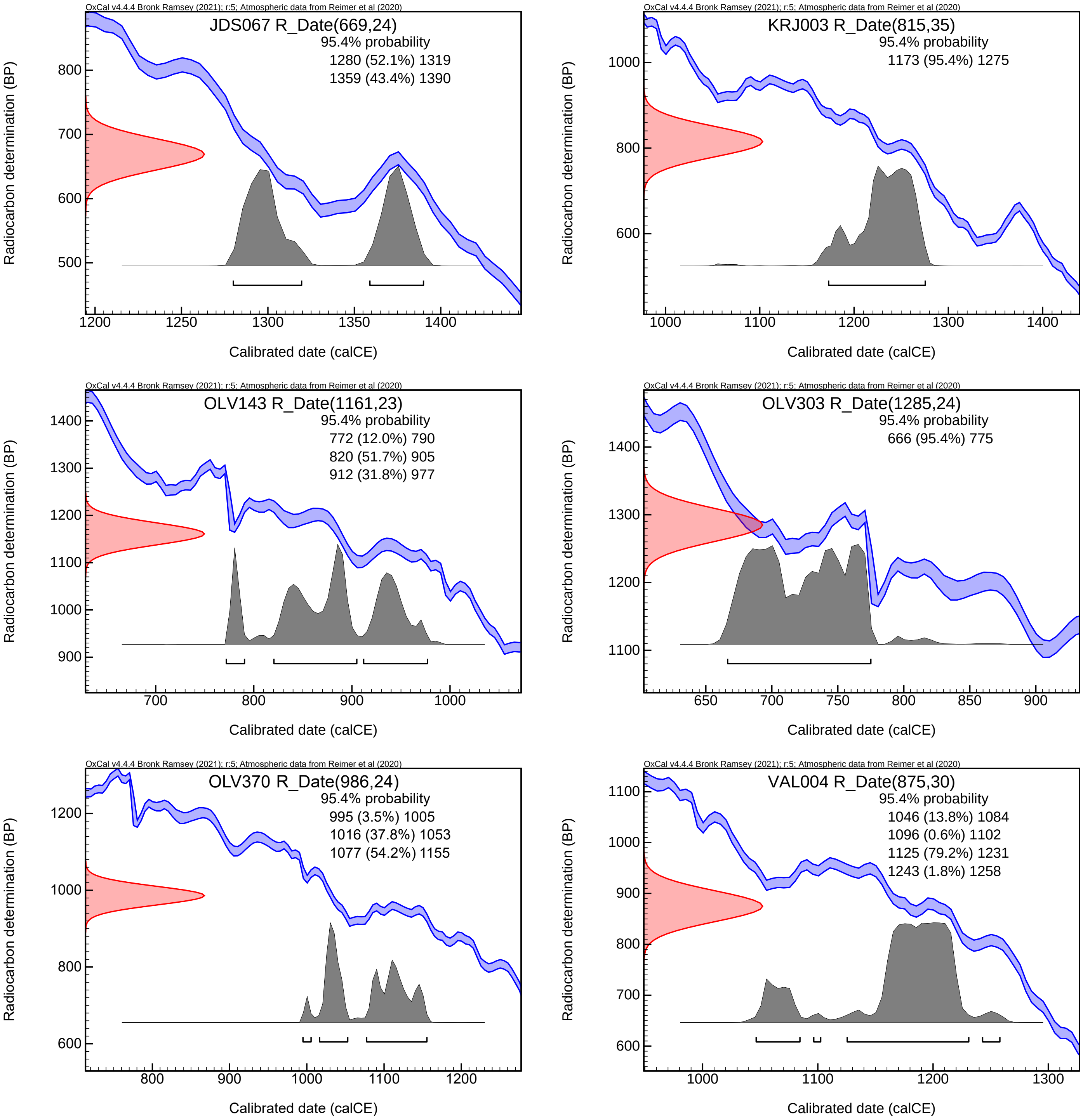


**Figure S5:** Calibration plots for radiocarbon dates for the individuals positive for HHV-6A, calibrated with IntCal20 in OxCal v4.4.4.

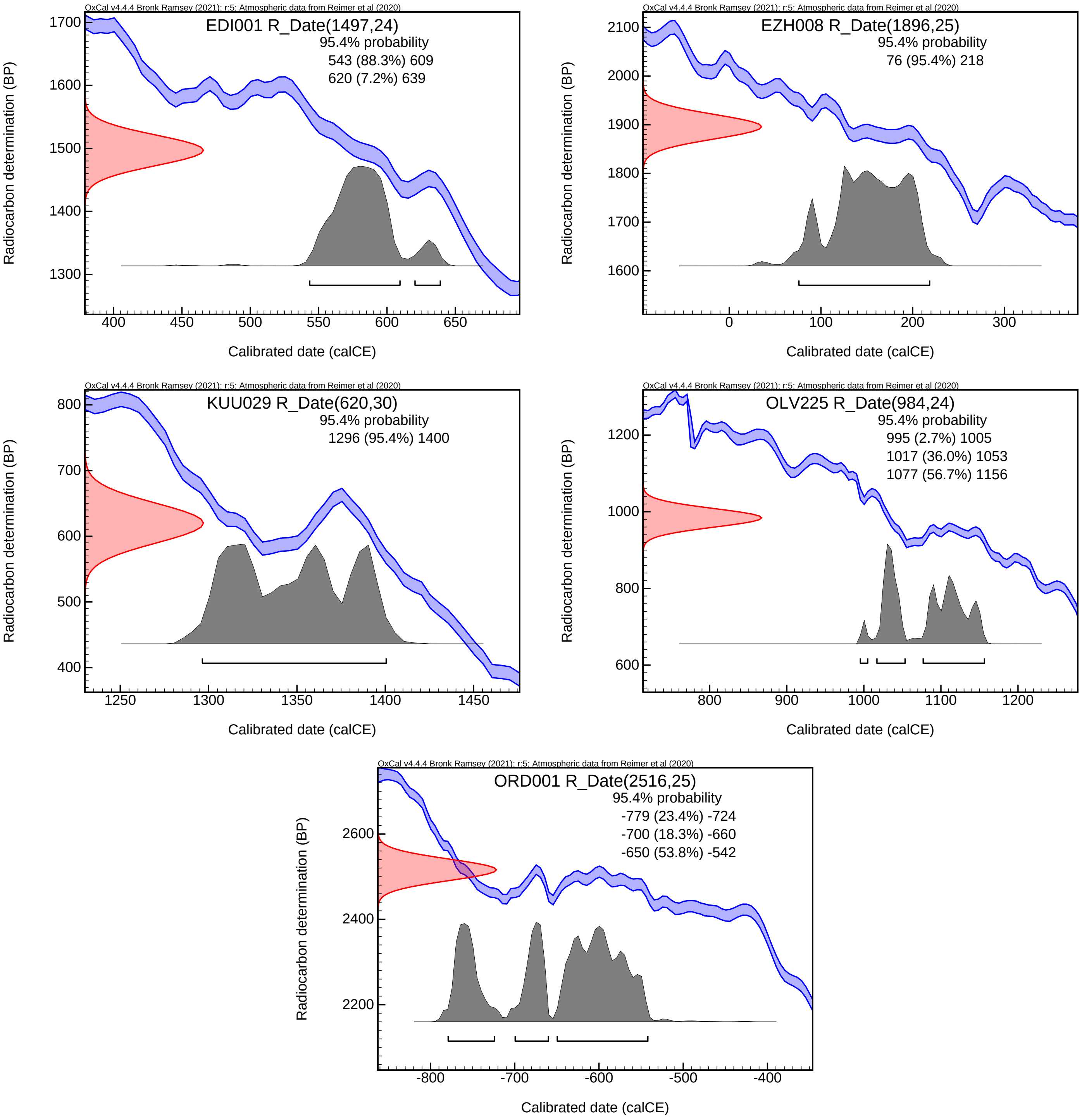


**Figure S6:** Calibration plots for radiocarbon dates for the individuals positive for HHV-6B, calibrated with IntCal20 in OxCal v4.4.4.

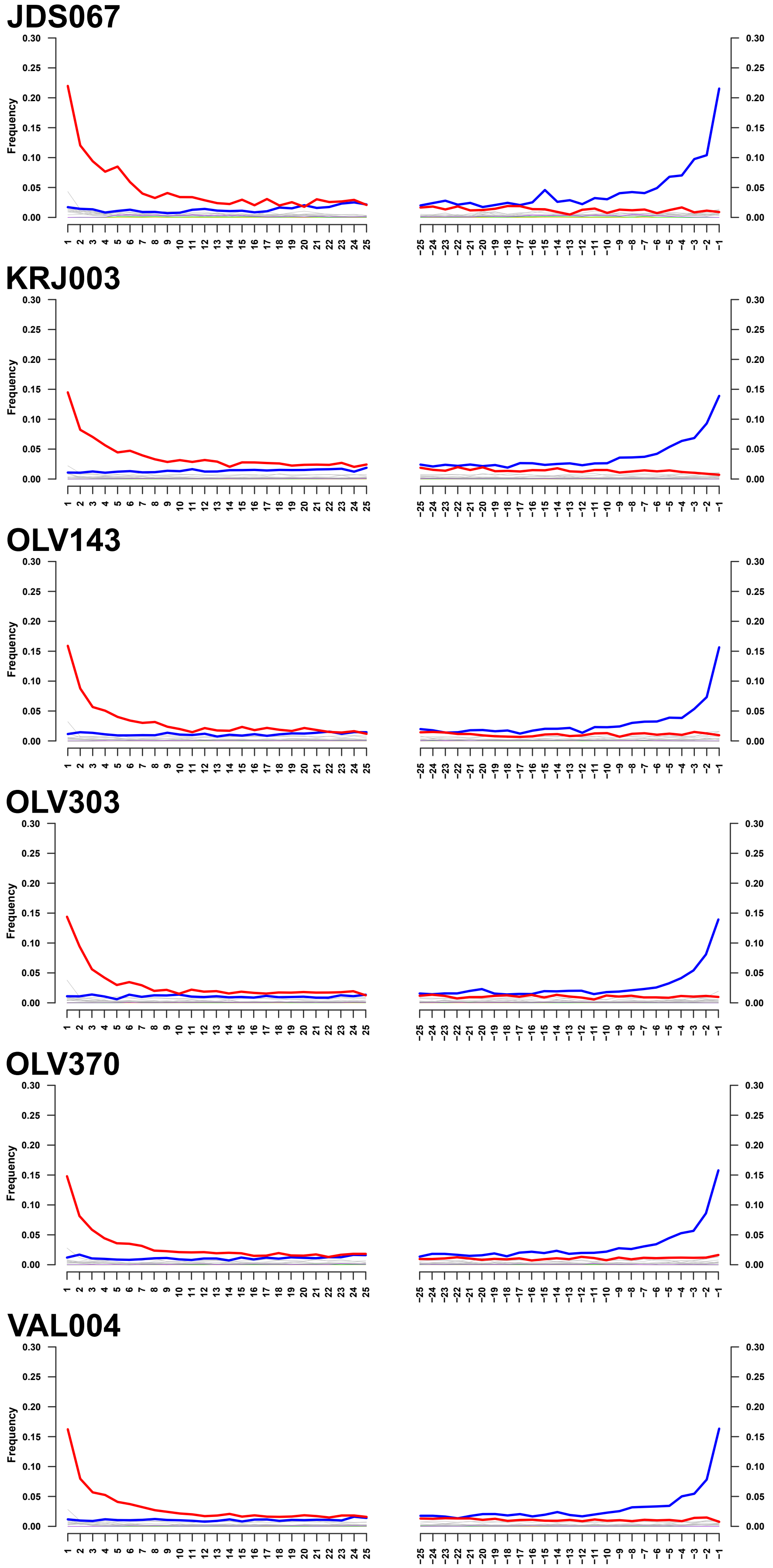


**Figure S7:** mapDamage2.0 deamination plots of our mappings to the HHV-6A reference sequence (NC_001664.4) for all HHV-6A positive samples.

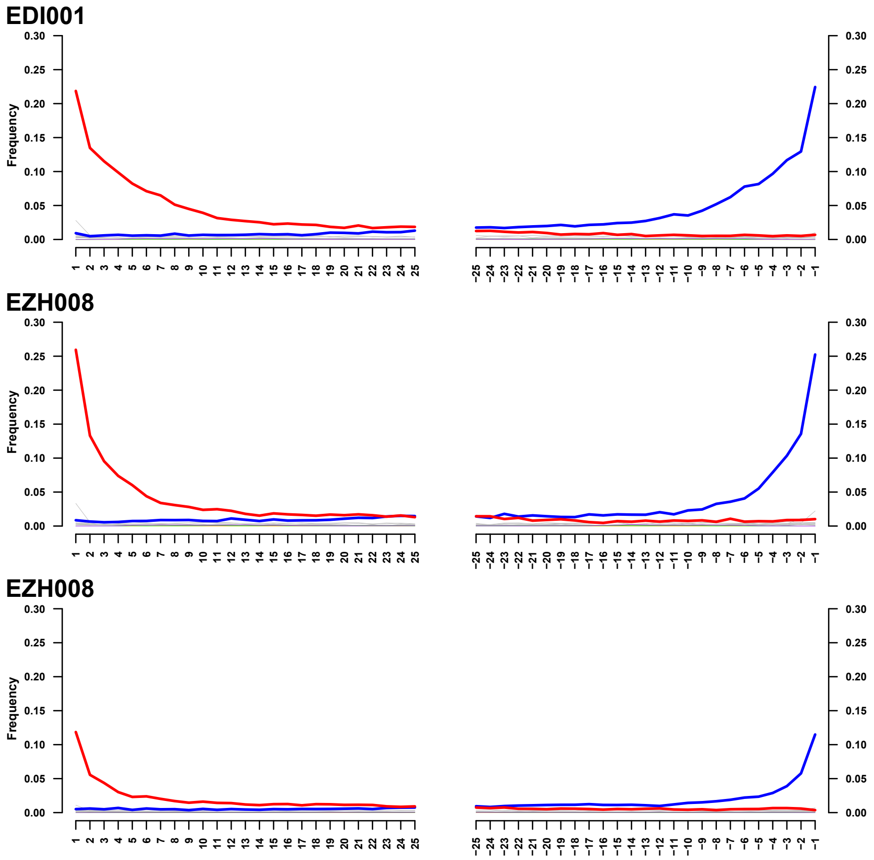


**Figure S8:** mapDamage2.0 deamination plots of our mappings to the HHV-6B reference sequence (NC_000898.1) for high coverage HHV-6B positive samples.

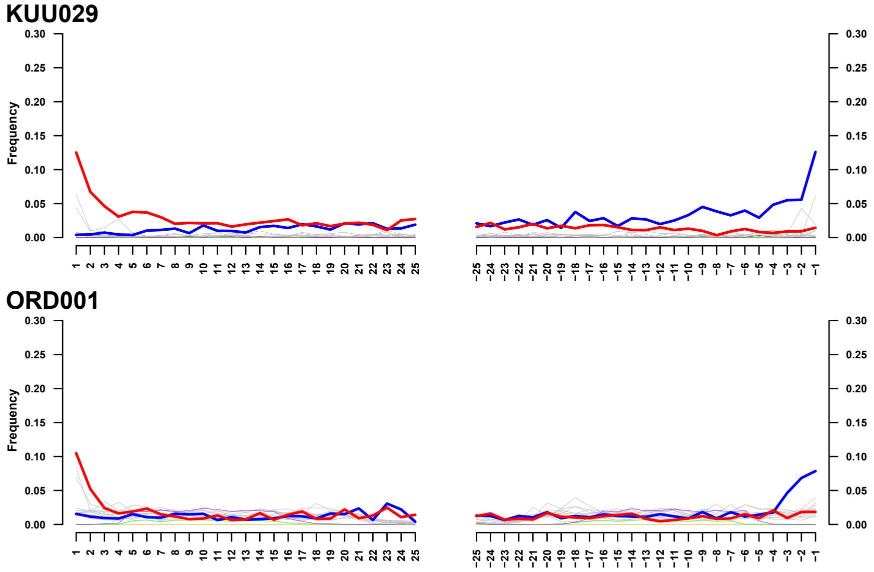


**Figure S9:** mapDamage2.0 deamination plots of our mappings to the HHV-6B reference sequence (NC_000898.1) for low coverage HHV-6B positive samples.

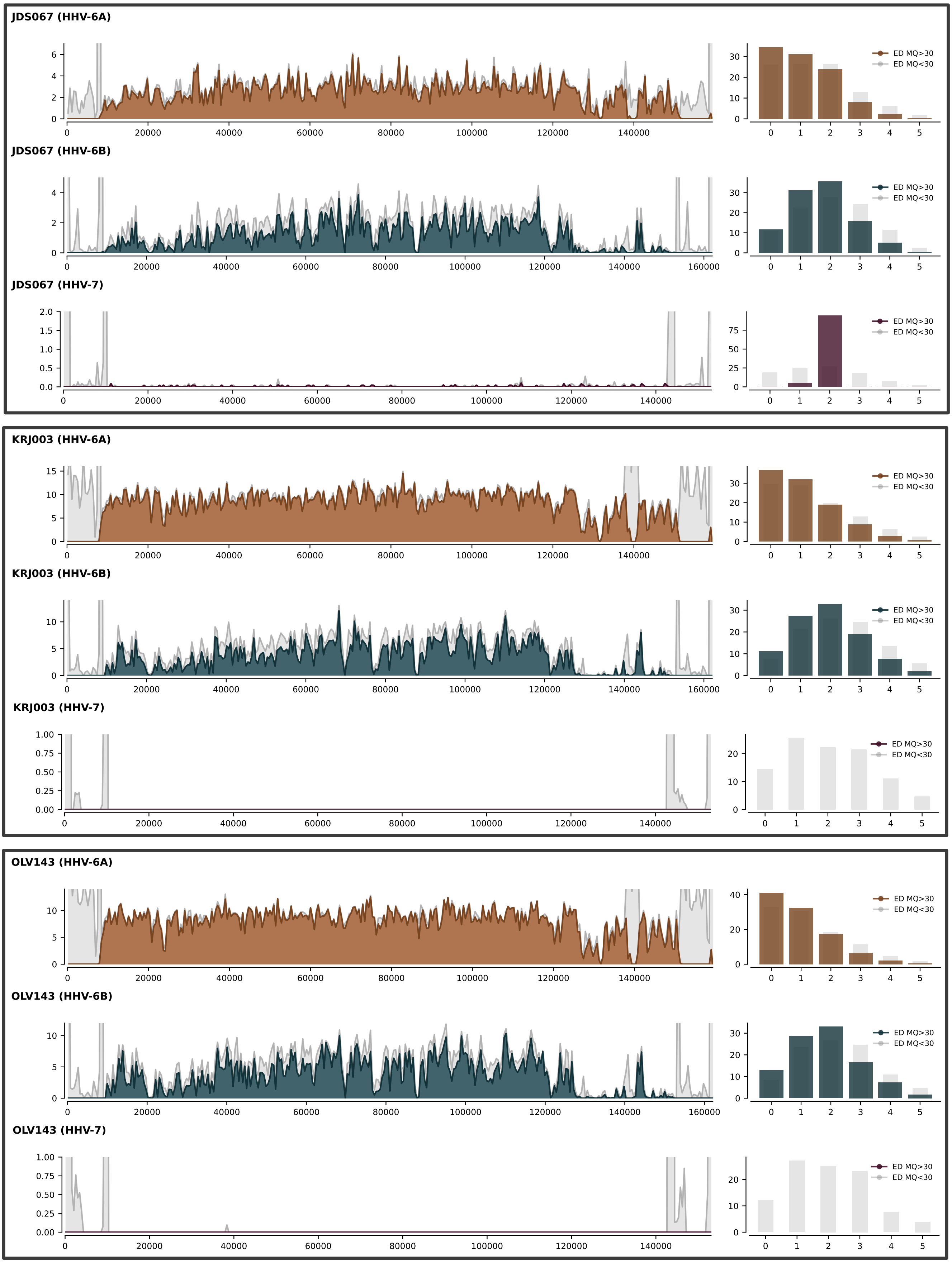


**Figure S10:** Coverage plots for all comparative mappings to respective reference sequences for HHV-6A (NC_001664.4) in ochre, HHV-6B (NC_000898.1) in dark blue and HHV-7 (NC_001716.2) in dark purple for HHV-6A positive samples. Intervals with reads under mapping quality of 30 are shown in light gray (depth of coverage shown on the y-axis and genome coordinates on the x-axis of the left plot). On the right, edit distances for the mappings are shown in a barplot with light gray bars showing the percentage of reads under a mapping quality of 30 for each edit distance and coloured bars showing the same for reads with mapping quality equal or above 30.

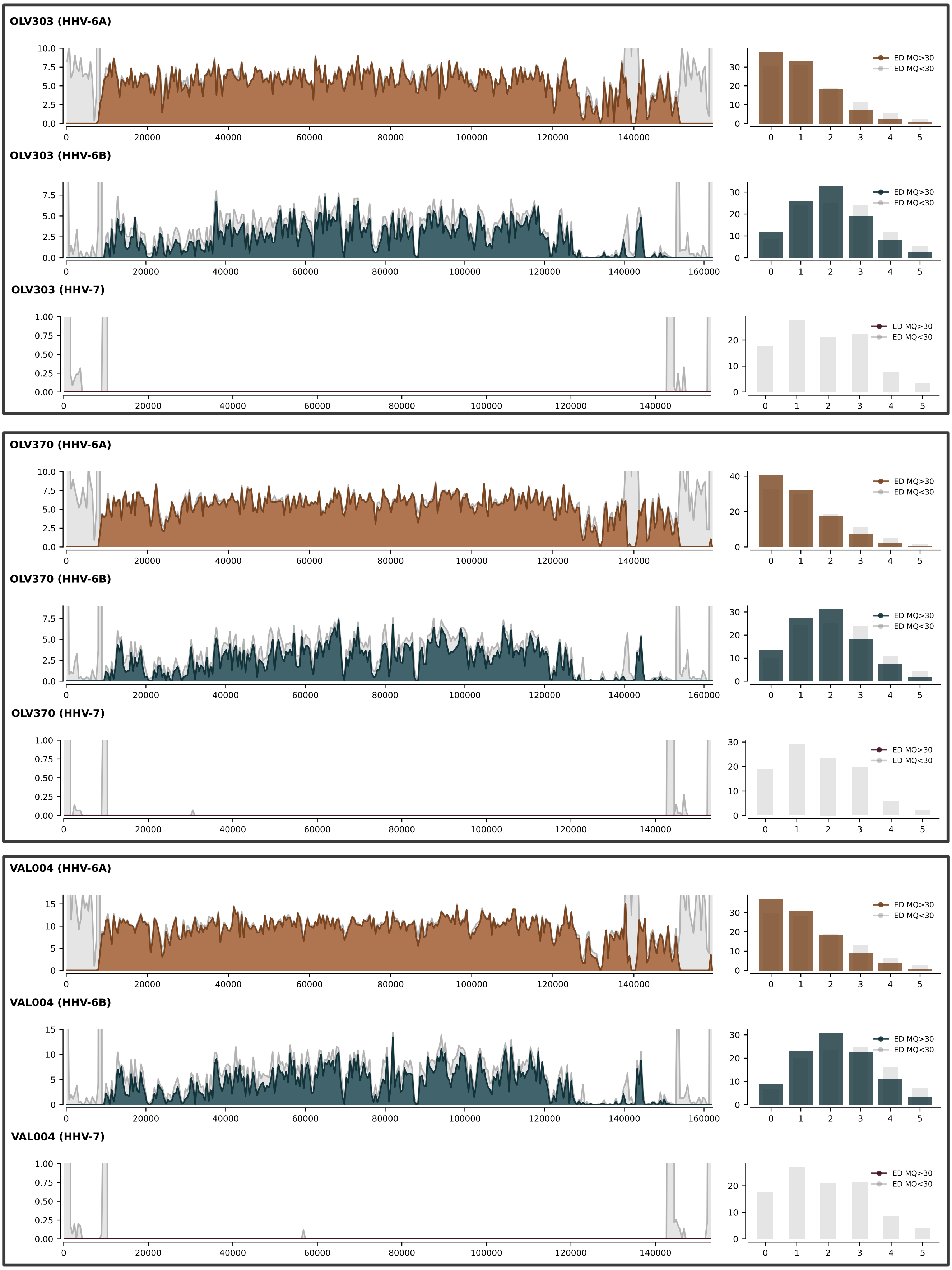


**Figure S11:** Coverage plots for all comparative mappings to respective reference sequences for HHV-6A (NC_001664.4) in ochre, HHV-6B (NC_000898.1) in dark blue and HHV-7 (NC_001716.2) in dark purple for HHV-6A positive samples. Intervals with reads under a mapping quality of 30 are shown in light gray (depth of coverage shown on the y-axis and genome coordinates on the x-axis of the left plot). On the right, edit distances for the mappings are shown in a barplot with light gray bars showing the percentage of reads under a mapping quality of 30 for each edit distance and coloured bars showing the same for reads with a mapping quality equal or above 30.

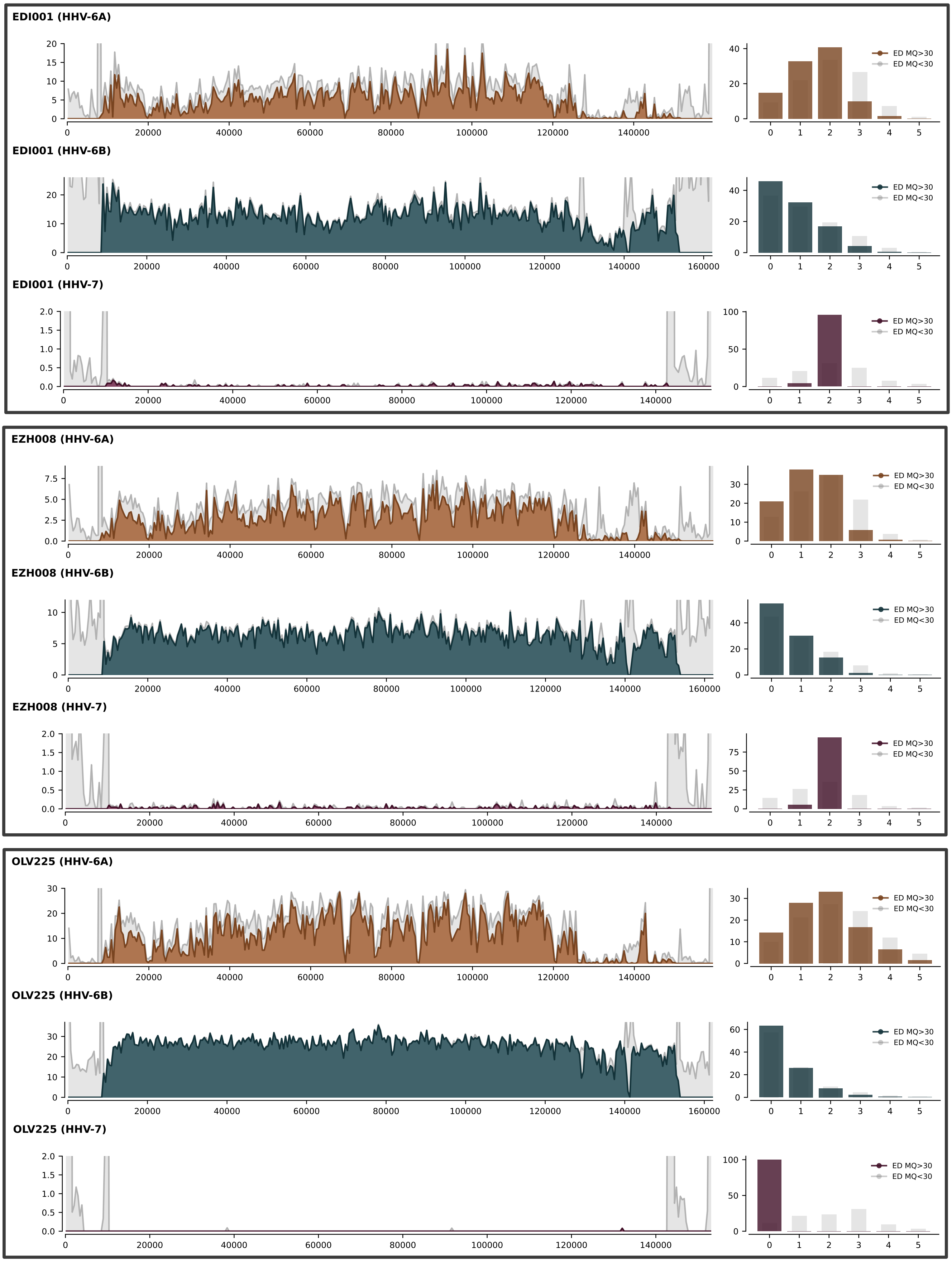


**Figure S12:** Coverage plots for all comparative mappings to respective reference sequences for HHV-6A (NC_001664.4) in ochre, HHV-6B (NC_000898.1) in dark blue and HHV-7 (NC_001716.2) in dark purple for all high coverage HHV-6B positive samples. Intervals with reads under a mapping quality of 30 are shown in light gray (depth of coverage shown on the y-axis and genome coordinates on the x-axis of the left plot). On the right, edit distances for the mappings are shown in a barplot with light gray bars showing the percentage of reads under a mapping quality of 30 for each edit distance and coloured bars showing the same for reads with a mapping quality equal or above 30.

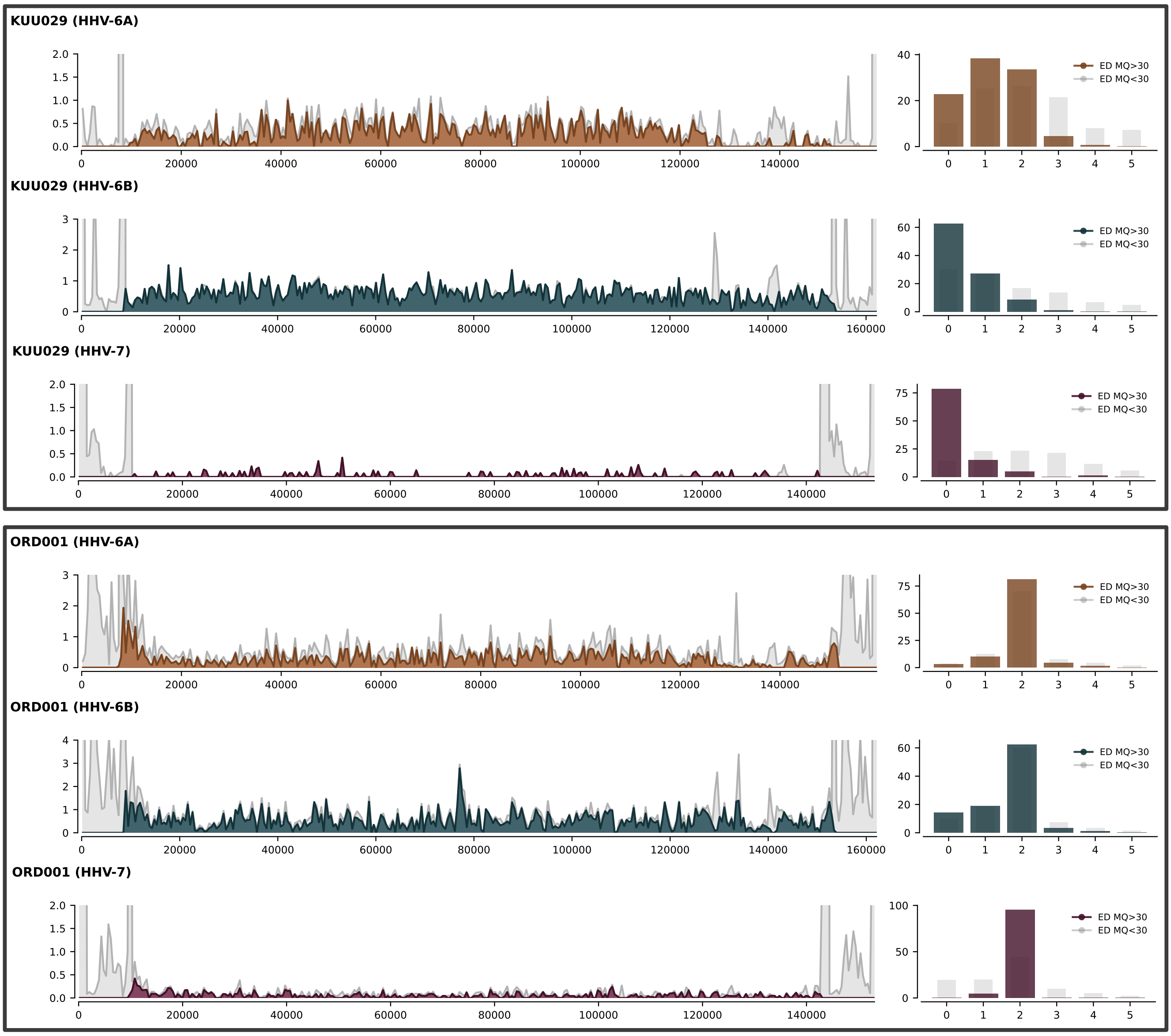


**Figure S13:** Coverage plots for all comparative mappings to respective reference sequences for HHV-6A (NC_001664.4) in ochre, HHV-6B (NC_000898.1) in dark blue and HHV-7 (NC_001716.2) in dark purple for all low coverage HHV-6B positive samples. Intervals with reads under a mapping quality of 30 are shown in light gray. On the right, edit distances for the mappings are shown in a barplot with light gray bars showing the percentage of reads under a mapping quality of 30 for each edit distance and coloured bars showing the same for reads with a mapping quality equal or above 30.

U12

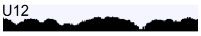

**Sample**: EDI001
**SNP**: 23328 T>G
**Variant Type**: stop_lost & splice_region_variant

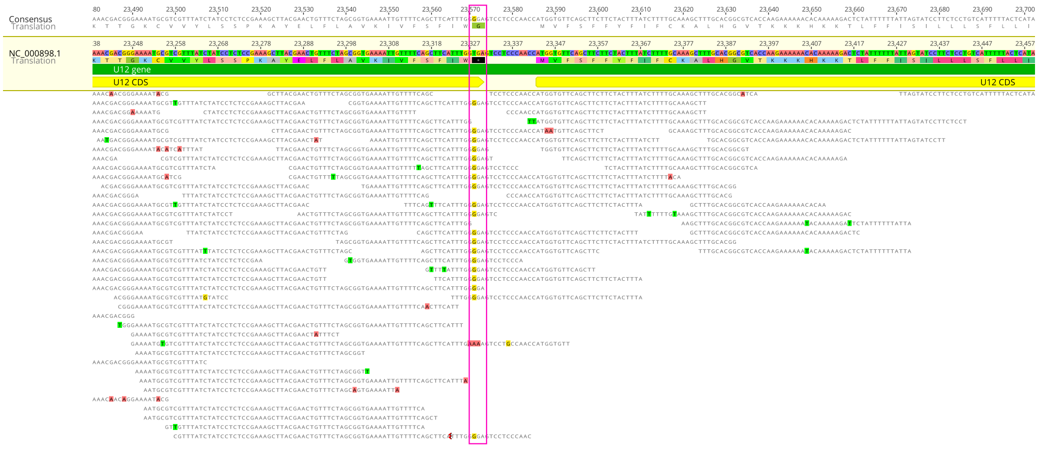


U67

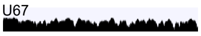

**Sample**: EDI001
**SNP**: 103758 G>A
**Variant Type**: start_lost

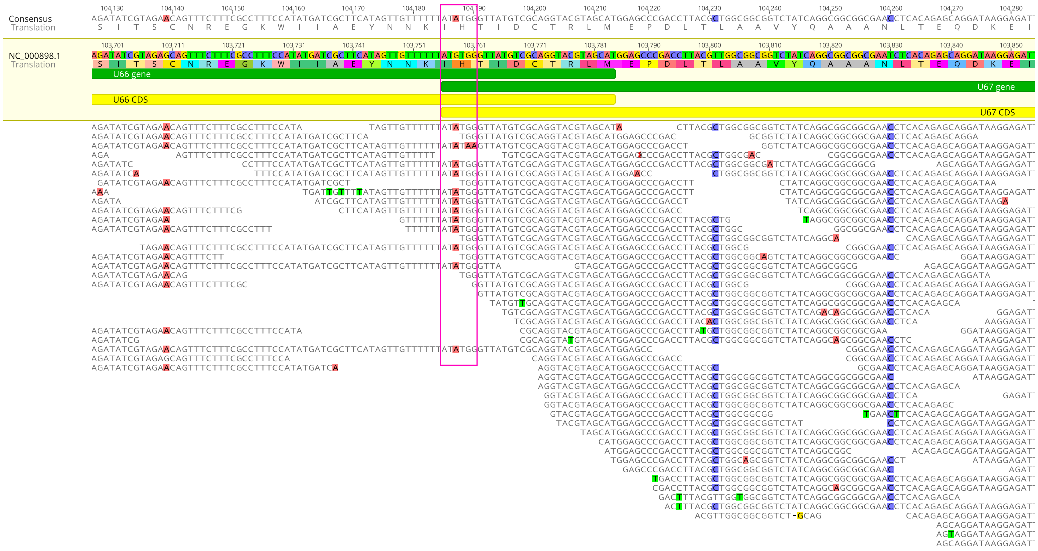


U91

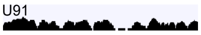

**Sample**: EZH008
**SNP**: 138935 GAA>GA
**Variant Type**: frameshift_variant

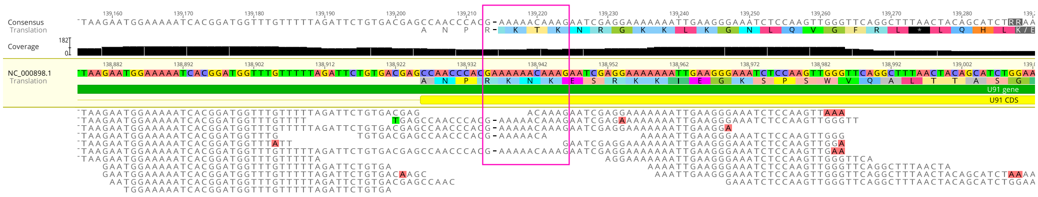


B9

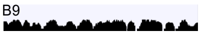

**Sample**: EZH008
**SNP**: 153242 CTT>CTTT
**Variant Type**: frameshift_variant

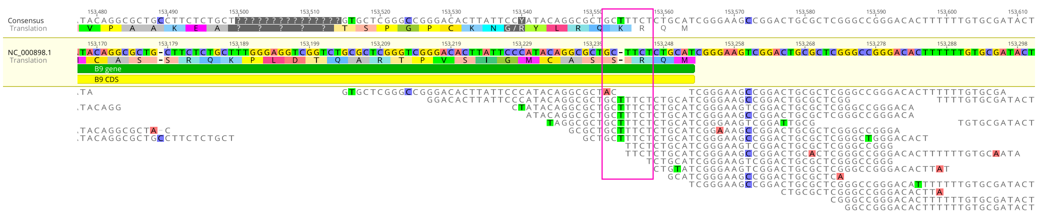


**Figure S14a:** Alignments within intervals of variants with high predicted impact by SNPEff, which fall within the unmasked regions of the genomes for the HHV-6B reference sequence. On the upper left is the log-scale coverage across the whole gene interval. SNP snapshots were generated with Geneious Prime 2023.2.1.

U12

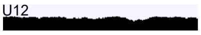

**Sample**: OLV225
**SNP**: 23328 T>G
**Variant Type**: stop_lost & splice_region_variant

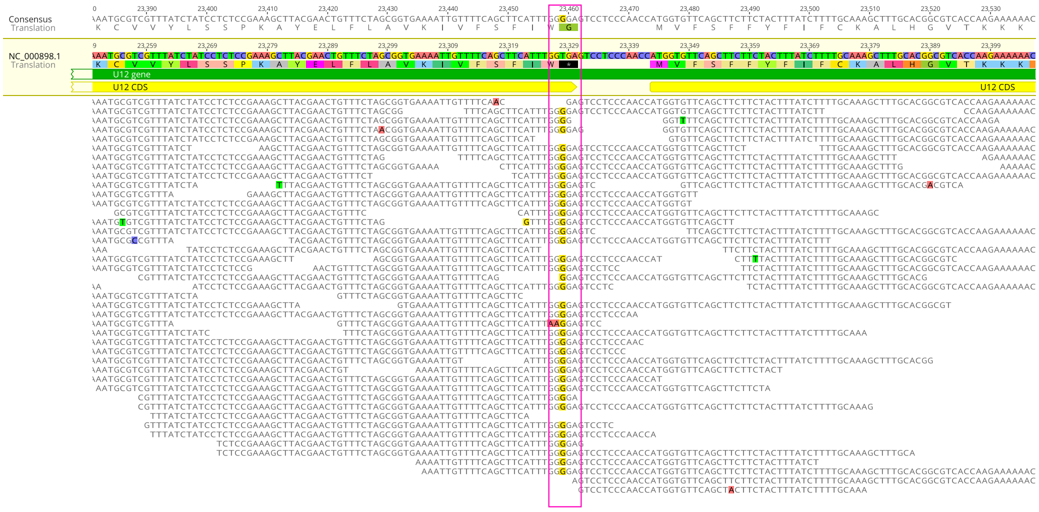


U67

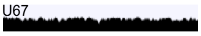

**Sample**: OLV225
**SNP**: 103758 G>A
**Variant Type**: start_lost

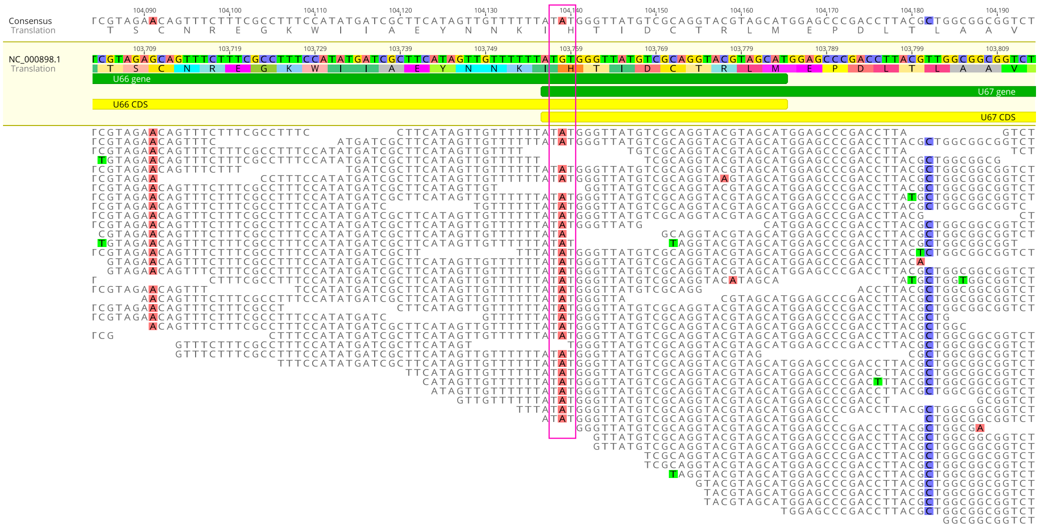


U95

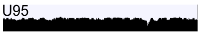

**Sample**: OLV225
**SNP**: 145618 C>T
**Variant Type**: stop_gained

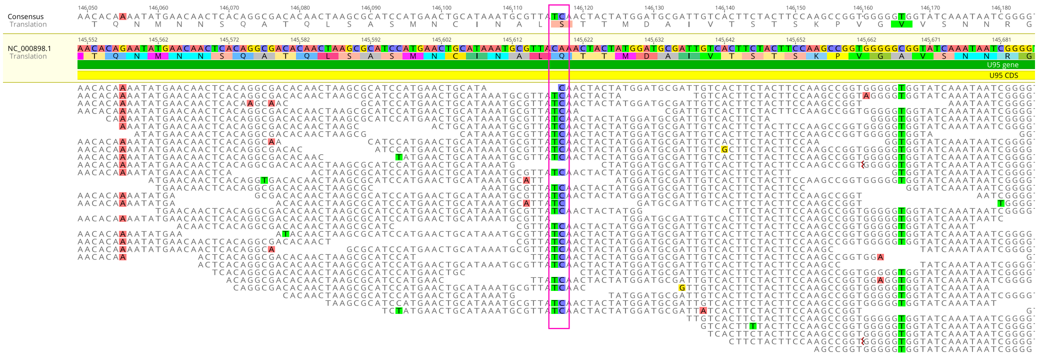


**Figure S14b:** Alignments within intervals of variants with high predicted impact by SNPEff, which fall within the unmasked regions of the genomes for the HHV-6B reference sequence. On the upper left is the log-scale coverage across the whole gene interval. SNP snapshots were generated with Geneious Prime 2023.2.1.

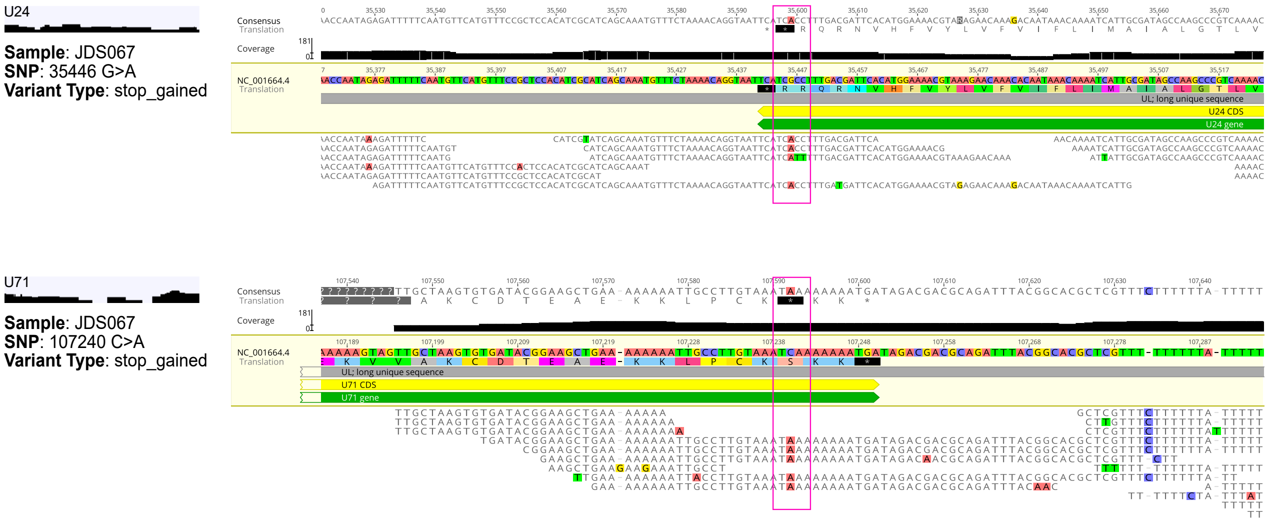


**Figure S14c:** Alignments within intervals of variants with high predicted impact by SNPEff, which fall within the unmasked regions of the genomes for the HHV-6A reference sequence. On the upper left is the log-scale coverage across the whole gene interval. SNP snapshots were generated with Geneious Prime 2023.2.1.

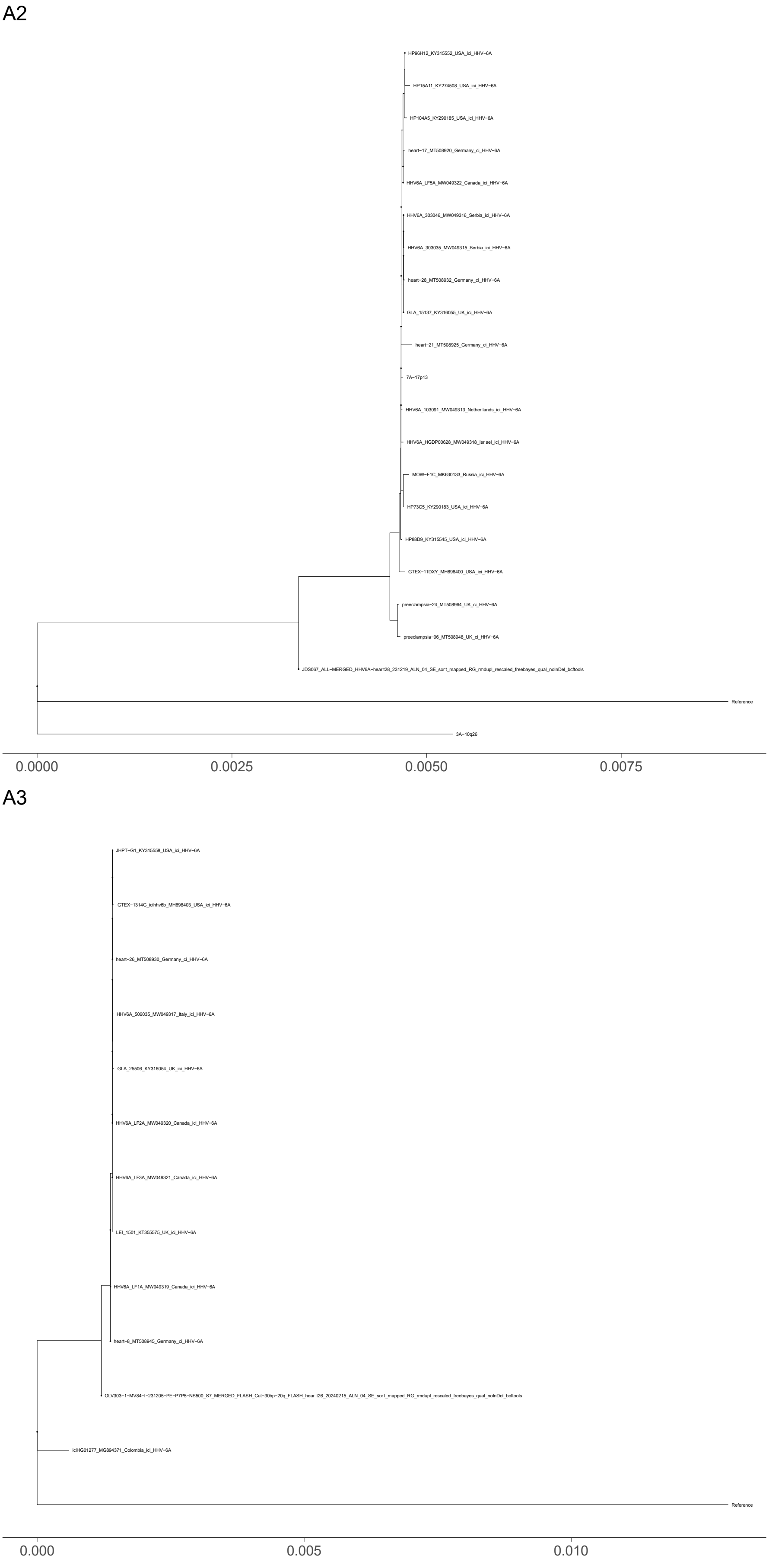


**Figure S15:** Maximum parsimony reconstruction of the homoplasy stripped core genome alignment for integrated clade A2 (top) and A3 (bottom).

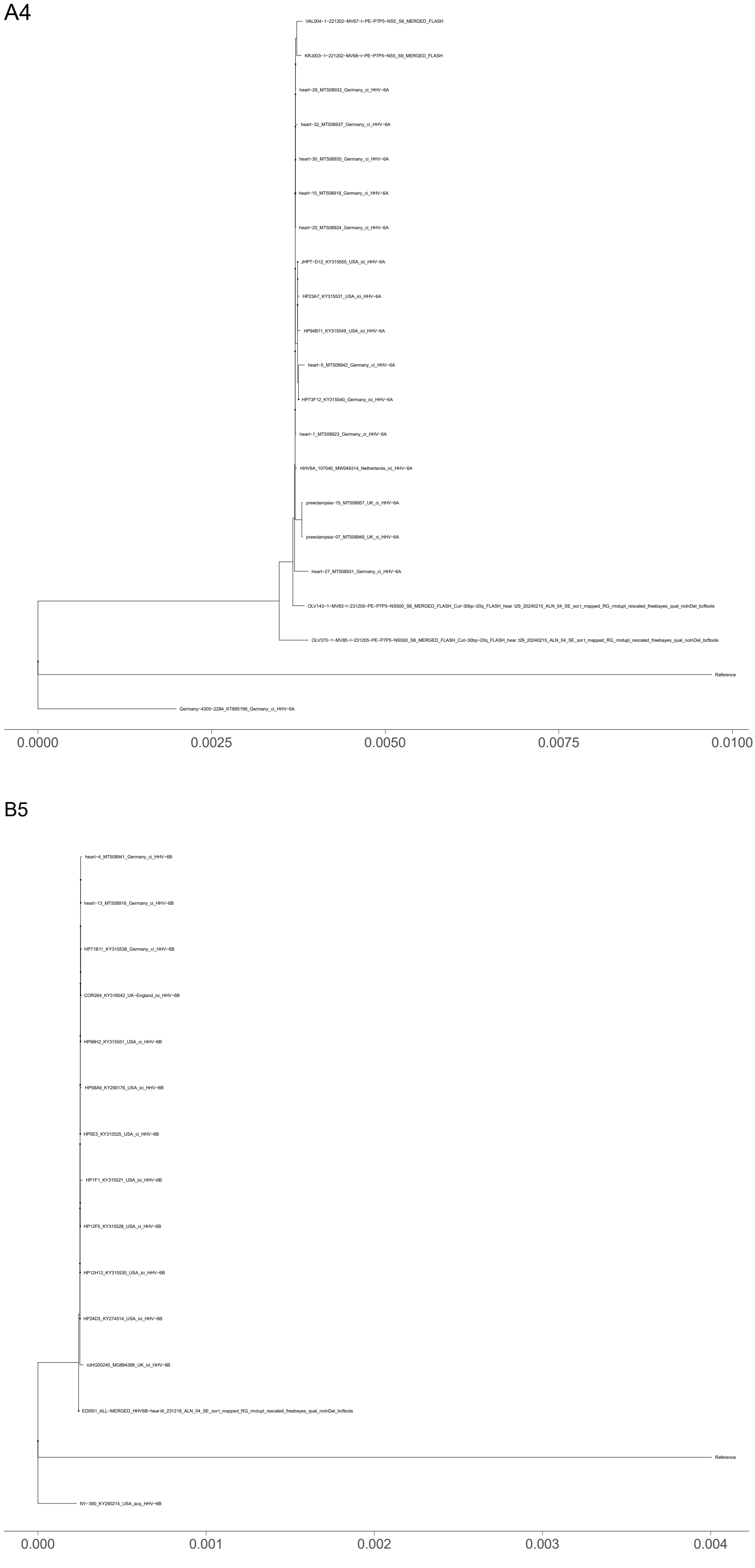


**Figure S16:** Maximum parsimony reconstruction of the homoplasy stripped core genome alignment for integrated clade A4 (top) and B5 (bottom).

B8

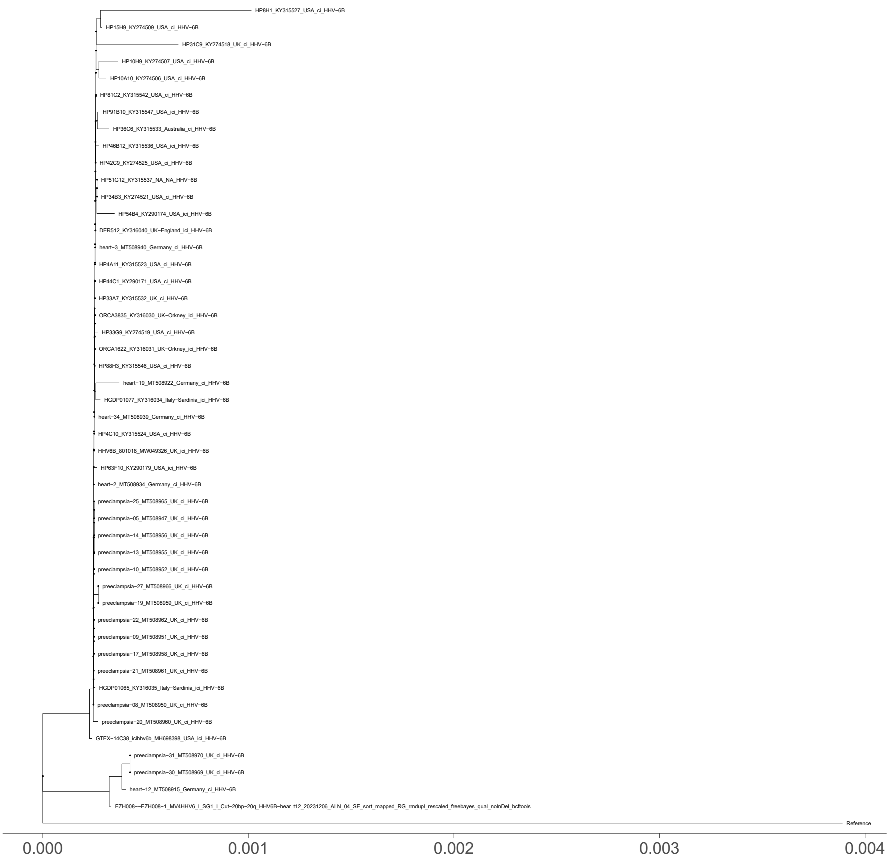


**Figure S17:** Maximum parsimony reconstruction of the homoplasy stripped core genome alignment for integrated clade B8.

### IX. SUPPLEMENTARY INFORMATION REFERENCES


1. De Winter, N. (2023). Eindverslag Sint-Truiden Groenmarkt. Opgraving naar aanleiding van de herinrichting van de Groenmarkt, het Trudoplein, de Diesterstraat, Plankstraat en Meinstraat. ARON rapport *1258*.

15. Masterson, E.E., Fitzpatrick, A.L., Enquobahrie, D.A., Mancl, L.A., Conde, E., and Hujoel, P.P. (2017). Malnutrition-related early childhood exposures and enamel defects in the permanent

dentition: A longitudinal study from the Bolivian Amazon. Am. J. Phys. Anthropol. *164*, 416–423. 10.1002/ajpa.23283.

21. Hillson, S. (2014). Tooth Development in Human Evolution and Bioarchaeology (Cambridge University Press).

22. Malim, T., and Hines, J. (1998). The Anglo-Saxon Cemetery at Edix Hill (Barrington A) (Council for British Archaeology).

23. Keller, M., Spyrou, M.A., Scheib, C.L., Neumann, G.U., Kröpelin, A., Haas-Gebhard, B., Päffgen, B., Haberstroh, J., Ribera I Lacomba, A., Raynaud, C., et al. (2019). Ancient Yersinia pestis genomes from across Western Europe reveal early diversification during the First Pandemic (541-750). Proc. Natl. Acad. Sci. U. S. A. *116*, 12363–12372. 10.1073/pnas.1820447116.

24. Guellil, M., Keller, M., Dittmar, J.M., Inskip, S.A., Cessford, C., Solnik, A., Kivisild, T., Metspalu, M., Robb, J.E., and Scheib, C.L. (2022). An invasive Haemophilus influenzae serotype b infection in an Anglo-Saxon plague victim. Genome Biol. *23*, 22. 10.1186/s13059-021-02580-z.

25. Lõhmus, M., Jonuks, T., and Malve, M. (2011). Archaeological salvage excavations at Kukruse: a Modern Age road, 13th–15th century cemetery. In Archaeological Fieldwork in Estonia 2010, E. Oras and E. Russow, eds. (Muinsuskaitseamet), pp. 103–114.

26. Malve, M. Kukruse matuste osteoloogiline analüüs (Unpublished Manuscript.).

27. Kustin, A. (1958). KALMISTU XIII—XIV SAJANDIST KARJAS, SAAREMAAL. Eesti NSV Teaduste Akadeemia Toimetised. Ühiskonnateaduste Seeria. 10.3176/hum.soc.sci.1958.1.05.

28. Mägi, M. (2002). At the Crossroads of Space and Time: Graves, Changing Society and Ideology on Saaremaa (Ösel), 9th-13th Centuries AD (Tallinn: Visby University Colleague - Institute of History).

29. Mägi, M., Malve, M., and Toome, T. (2018). Early Christian burials at Valjala churchyard, Saaremaa. Archaeological Fieldwork in Estonia, 98–118.

30. Turkina, T.Y. (2018). Ezhol burial ground of the 5th–6th centuries AD on the Middle Vychegda (Initial research results). In Матвеевой и 70-летию со дня рождения И.Б. Васильева : Материалы Всероссийской археологической конференции с международным участием 8-11 октября 2018 (Самара).

31. Silaev VI, Belitskaya AL, Turkina TYu., Smoleva IV, Khazov AF, Kiseleva D (2019). Environment and diet of the Early Medieval population of the European Northeast (according to isotopic-geochemical analysis of the anthropological materials from burial grounds of the V-VII centuries A.D.). Известия Коми научного центра УРО РАН, 53–64.

32. Bronk Ramsey, C. (2021). OxCal 4.4 Manual.

33. Reimer, P.J., Austin, W.E.N., Bard, E., Bayliss, A., Blackwell, P.G., Ramsey, C.B., Butzin, M., Cheng, H., Lawrence Edwards, R., Friedrich, M., et al. (2020). The IntCal20 Northern Hemisphere Radiocarbon Age Calibration Curve (0–55 cal kBP). Radiocarbon *62*, 725–757. 10.1017/RDC.2020.41.

34. Iker, R. (1986). Ordona VII/2: Les tombes dauniennes: les tombes du IVe et du début du IIIe siècles avant notre ère. Preprint at Brussels: Academia Belgica.

35. Scaggion, C., and Carrara, N. (2016). New studies on human skeletal remains from the ancient Her-donia ( southeast Italy ) . Evidences of tuberculosis and brucellosis : two diseases connected with farm animals. Antrocom: Online Journal of Anthropology.

36. Aneli, S., Saupe, T., Montinaro, F., Solnik, A., Molinaro, L., Scaggion, C., Carrara, N., Raveane, A., Kivisild, T., Metspalu, M., et al. (2022). The Genetic Origin of Daunians and the Pan-Mediterranean Southern Italian Iron Age Context. Mol. Biol. Evol. *39*. 10.1093/molbev/msac014.

37. Pockrandt, C., Alzamel, M., Iliopoulos, C.S., and Reinert, K. (2020). GenMap: ultra-fast computation of genome mappability. Bioinformatics *36*, 3687–3692. 10.1093/bioinformatics/btaa222.

38. Martin, D.P., Varsani, A., Roumagnac, P., Botha, G., Maslamoney, S., Schwab, T., Kelz, Z., Kumar, V., and Murrell, B. (2021). RDP5: a computer program for analyzing recombination in, and removing signals of recombination from, nucleotide sequence datasets. Virus Evol *7*, veaa087. 10.1093/ve/veaa087.

39. Zhang, E., Bell, A.J., Wilkie, G.S., Suárez, N.M., Batini, C., Veal, C.D., Armendáriz-Castillo, I., Neumann, R., Cotton, V.E., Huang, Y., et al. (2017). Inherited Chromosomally Integrated Human Herpesvirus 6 Genomes Are Ancient, Intact, and Potentially Able To Reactivate from Telomeres. J. Virol. *91*. 10.1128/JVI.01137-17.

40. Finkel, Y., Schmiedel, D., Tai-Schmiedel, J., Nachshon, A., Winkler, R., Dobesova, M., Schwartz, M., Mandelboim, O., and Stern-Ginossar, N. (2020). Comprehensive annotations of human herpesvirus 6A and 6B genomes reveal novel and conserved genomic features. Elife *9*. 10.7554/eLife.50960.

41. Greninger, A.L., Knudsen, G.M., Roychoudhury, P., Hanson, D.J., Sedlak, R.H., Xie, H., Guan, J., Nguyen, T., Peddu, V., Boeckh, M., et al. (2018). Comparative genomic, transcriptomic, and proteomic reannotation of human herpesvirus 6. BMC Genomics *19*, 204. 10.1186/s12864-018-4604-2.

42. Touraine, J.L., Betuel, H., Souillet, G., and Jeune, M. (1978). Combined immunodeficiency disease associated with absence of cell-surface HLA-A and -B antigens. J. Pediatr. *93*, 47–51.

10.1016/s0022-3476(78)80598-8.

43. McKenna, A., Hanna, M., Banks, E., Sivachenko, A., Cibulskis, K., Kernytsky, A., Garimella, K., Altshuler, D., Gabriel, S., Daly, M., et al. (2010). The Genome Analysis Toolkit: a MapReduce framework for analyzing next-generation DNA sequencing data. Genome Res. *20*, 1297–1303. 10.1101/gr.107524.110.

44. Lam, H.M., Ratmann, O., and Boni, M.F. (2018). Improved Algorithmic Complexity for the 3SEQ Recombination Detection Algorithm. Mol. Biol. Evol. *35*, 247–251. 10.1093/molbev/msx263.

45. Minh, B.Q., Schmidt, H.A., Chernomor, O., Schrempf, D., Woodhams, M.D., von Haeseler, A., and Lanfear, R. (2020). IQ-TREE 2: New Models and Efficient Methods for Phylogenetic Inference in the Genomic Era. Mol. Biol. Evol. *37*, 1530–1534. 10.1093/molbev/msaa015.

46. Didelot, X., Croucher, N.J., Bentley, S.D., Harris, S.R., and Wilson, D.J. (2018). Bayesian inference of ancestral dates on bacterial phylogenetic trees. Nucleic Acids Res. *46*, e134. 10.1093/nar/gky783.

47. Hoang, D.T., Vinh, L.S., Flouri, T., Stamatakis, A., von Haeseler, A., and Minh, B.Q. (2018). MPBoot: fast phylogenetic maximum parsimony tree inference and bootstrap approximation. BMC Evol. Biol. *18*, 11. 10.1186/s12862-018-1131-3.

48. Crispell, J., Balaz, D., and Gordon, S.V. (2019). HomoplasyFinder: a simple tool to identify homoplasies on a phylogeny. Microb Genom *5*. 10.1099/mgen.0.000245.

49. Aswad, A., Aimola, G., Wight, D., Roychoudhury, P., Zimmermann, C., Hill, J., Lassner, D., Xie, H., Huang, M.-L., Parrish, N.F., et al. (2021). Evolutionary History of Endogenous Human Herpesvirus 6 Reflects Human Migration out of Africa. Mol. Biol. Evol. *38*, 96–107. 10.1093/molbev/msaa190.
